# Crystal structure of calcium bound outer membrane phospholipase A (OmpLA) from *Salmonella typhi* and *in silico* anti-microbial screening

**DOI:** 10.1101/2020.01.08.898262

**Authors:** Perumal Perumal, Rahul Raina, Sundara Baalaji Narayanan, Arulandu Arockiasamy

## Abstract

Antimicrobial resistance is widespread in *Salmonella* infections that affect millions worldwide. *Salmonella typhi* and other Gram-negative bacterial pathogens encode an outer membrane phospholipase A (OmpLA), crucial for their membrane integrity. Further, OmpLA is implicated in pathogen internalization, haemolysis, acid tolerance, virulence and sustained infection in human hosts. OmpLA is an attractive drug target for developing novel anti-microbials that attenuate virulence, as the abrogation of OmpLA encoding *pldA* gene causes loss of virulence. Here, we present the crystal structure of *Salmonella typhi* OmpLA in dimeric calcium bound activated state at 2.95 Å. Structure analysis suggests that OmpLA is a potential druggable target. Further, we have identified and shortlisted small molecules that bind at the dimer interface using structure based *in silico* screening, docking and molecular dynamics. While it requires further experimental validation, anti-microbial discovery targeting OmpLA from gram-negative pathogens offers an advantage as OmpLA is required for virulence.

## Introduction

*Salmonella typhi*, a human pathogen, causes typhoid fever that affects ∼ 21 million people every year (WHO, Fact sheet 2018). Existing therapies include various antibiotics, misuse of which results in rampant antimicrobial resistance. The problem of emergence of resistant strains, in part, is due to antibiotics targeting essential genes and pathways. Thus, virulence causing factors are an attractive and alternate molecular target to design novel antimicrobials. Bacterial outer membrane proteins are involved in signal transduction and transport of nutrients with few acting as enzymes, one of which is outer membrane phospholipase A (OmpLA) encoded by *pldA*. OmpLA *encoding pldA* from *Escherichia coli, Salmonella typhimurium, Klebsiella pneumoniae*, and *Proteus Vulgaris* were extensively explored for its function^1,2^. OmpLA is a highly conserved protein essential for bacterial membrane integrity and is present in all members of the Enterobacteriaceae family. OmpLA shows enzymatic activity similar to those of soluble phospholipases A1 and A2 as well as that of 1-acyl- and 2-acyl-lysophospholipase and diacylglyceride lipase^3^. *E. coli* OmpLA (EcOmpLA) is shown to play key role during secretion of bacteriocins^4,5^. Though functionally inactive during normal growth phase^6^, OmpLA shows increased enzymatic activity during membrane damage, triggered by phage-mediated lysis^7^ or temperature shock^8^. OmpLA mutant of *Shigella flexneri* shows altered expression of membrane-integrated proteins and affects expression of ABC transporters and type III secretion system function^9^. Further, OmpLA is also implicated in various bacterial pathologies such as massive tissue destruction related to gas gangrene, sepsis, skin and lung infections^10^. Thus, the existing data strongly suggests OmpLA is not essential for growth but is a major virulence factor and hence a potential drug target. Interestingly, bacterial OmpLA shows no sequence or structural homology with soluble phospholipases in human, indicative of its usefulness as a unique drug target.

## Results and discussion

### Structure determination of *S. typhi* OmpLA

*S. typhi* OmpLA (StOmpLA), cloned without the signal sequence (Fig**. S**1a, b), was overexpressed in *E. coli* as inclusion bodies. Scouting urea concentration for unfolding OmpLA shows sharp rise in absorption at OD_280_ at 4 M which stabilizes at 6 M (Fig. S1c, d). Large-scale unfolding was done using 8 M urea, and refolding was achieved in presence of 0.3% Polyoxyethylene (9) lauryl ether (C_12_E_9)_. Refolded OmpLA was further purified using ion-exchange and size exclusion chromatography (Fig. S1e, f) and concentrated to 14 mg/ml. Then the protein was detergent exchanged from C_12_E_9_ to β-octyl-glucoside (β-OG) using an ion-exchange column, concentrated to 14 mg/ml, flash frozen in liquid nitrogen and stored at −80 °C before crystallization. Total yield of refolded and detergent exchanged OmpLA was 15 mg/gm of purified inclusion bodies. Circular dichroism (CD) data (Fig. S1g), using a 240 to 205 nm scan, shows minimum mean residue molecular ellipticity [Θ] (MRW) around 217 nm and a crossover at 210 nm, indicating refolded OmpLA contains mainly of β-structure, similar to EcOmpLA^11,12,13^. Protocol described here (Methods) yields more refolded OmpLA than previously published methods^14,15^. Refolded OmpLA was crystallized in various conditions using MemGold 1 and 2. Diffraction quality crystals (Fig. S1h) were obtained in a condition containing 0.1 M sodium iodide, 0.1 M sodium phosphate (pH 7.0), and 33% v/v polyethylene glycol 300. StOmpLA crystals were directly mounted on home source X-ray and screened for diffraction quality. A single crystal diffracted to 2.95 Å resolution, belonging to the space group P2_1_2_1_2_1_ with the unit cell parameters; a=79.340Å, b=83.389Å, c=95.463Å and α = β = γ = 90°. Mathew’s coefficient calculation suggested 48.63% solvent content and a Vm of 2.39 Å^3^/Dalton, indicating two molecules are present in the asymmetric unit. Molecular replacement using EcOmpLA (PDB: 1QD6) as template structure yielded initial phases. StOmpLA model was built, using COOT^16^, and refined to a final R and R_free_ of 23.8 and 28.9 (Table 1), and deposited to the Protein Data Bank with accession code 5DQX.

**Table 1.** Data collection and refinement statistics for StOmpLA (PDB:5DQX)

|  |  |
| --- | --- |
| Wavelength Å | 1.5418 |
| Resolution range Å | 34.43 - 2.95 (3.055 - 2.95) |
| Space group | P 21 21 21 |
| Unit cell | 79.340, 83.389, 95.463 Å, 90, 90, 90° |
| No. of images collected | 155 |
| Unique reflections | 13771 (1346) |
| Multiplicity | 2.8 |
| Completeness (%) | 99.32 (99.41) |
| Mean I/sigma(I) | 8.54 (2.63) |
| Wilson B-factor Å<sup>2</sup> | 36.33 |
| R-merge | 1.097e-17 (1.118e-17) |
| R-meas | 1.551e-17 (1.581e-17) |
| R-pim | 1.097e-17 (1.118e-17) |
| CC1/2 | 1 (1) |
| CC* | 1 (1) |
| Reflections used in refinement | 13765 (1346) |
| Reflections used for R-free | 685 (67) |
| R-work | 0.235 (0.238) |
| R-free | 0.282 (0.289) |
| CC (work) | 0.897 (0.738) |
| CC (free) | 0.955 (0.613) |
| **Number of non-hydrogen atoms** | 4099 |
| macromolecule | 4018 |
| ligands | 63 |
| solvent | 18 |
| Protein residues | 504 |
| RMS (bonds) Å<sup>2</sup> | 0.010 |
| RMS (angles) ° | 1.44 |
| Ramachandran favored (%) | 92.60 |
| Ramachandran allowed (%) | 6.00 |
| Ramachandran outliers (%) | 1.40 |
| Rotamer outliers (%) | 7.00 |
| Clash score | 6.72 |
| **Average B-factor Å<sup>2</sup>** | 29.36 |
| Macromolecule | 29.28 |
| Ligands | 35.14 |
| Solvent | 26.08 |

Statistics for the highest-resolution shell are shown in parentheses.

### Crystal structure of calcium bound *S. typhi* OmpLA dimer

StOmpLA is crystallized as a calcium bound homodimer with each monomer forming a β-barrel, containing two flat surfaces, facing the membrane bilayer, and two highly convex sides (Fig. 1a, b); one facing the periplasm and the other towards cytoplasm, respectively. Each β-barrel is comprised of 13 anti-parallel β-strands (β1 – β13), an α-helix (α1) (between β8 and β9) and three 3_10_ helices (η1-η3) with η1/η2 located between β2 and β3, and η3 located between β4 and β5. 3_10_ helices η1 and η2 form a helix-turn-helix motif towards the extracellular end of the β-barrel (Fig. 1c). The structure has 18 turns containing 2 α-turns (TTT), 8 β-turns (TT) and 8 long loops facing polar compartments. The loops between β-strands β2 and β3, β12 and β13 along with η1/η2 helix-turn-helix motif constrict the opening of barrel towards extracellular space, and N- and C-terminal loops cover the periplasmic ends of β-barrel (Fig. 1c). Temperature factor (B-factor) ranges between 16 to 74 Å^2^ with an average value of 29.36 Å^2^. The loop region preceding β1 strand, has a high B-factor (D46: 77 Å^2^, N47: 73 Å^2^, P48: 70 Å^2^) as shown in Figure 1d, e. The differences in B-factor of individual residues in each chain as shown in Figure 1e are not significant. Each monomer has two highly ordered aromatic belts (Fig. 2); one near extracellular space of the β-barrel and another near periplasmic space. Interaction between Y211 and Y272 brings loop 17 closer to the β-barrel thereby constricting the pore size (Fig. 2a). There are two sulphur-π pairs, present towards the interior of OmpLA channel formed by the residues M284 & W258 (4.5 Å), and M212 & W175 (6.0 Å) as shown in Figure 2b,c, help stabilize β13, α1, L12 and L13 with respect to the barrel, and thereby further constricting the channel opening towards the extracellular compartment. Superposition of StOmpLA monomers on Cα atoms shows RMSD of 0.096Å, suggesting no major structural differences between them. The minor differences observed are mainly confined to loops; L1 (Glu45-Thr51), L9 (Phe148-Trp151), L16 (Pro249 – Leu254) and residues in β5 strand (Fig. S2a), exposed to the periplasmic region. Loop 1 (70 Å^2^) and 9 (52 Å^2^) have a higher b-factor in comparison with the average b-factor of 29.3 Å^2^ (Fig. S2b).

**Figure 1.**
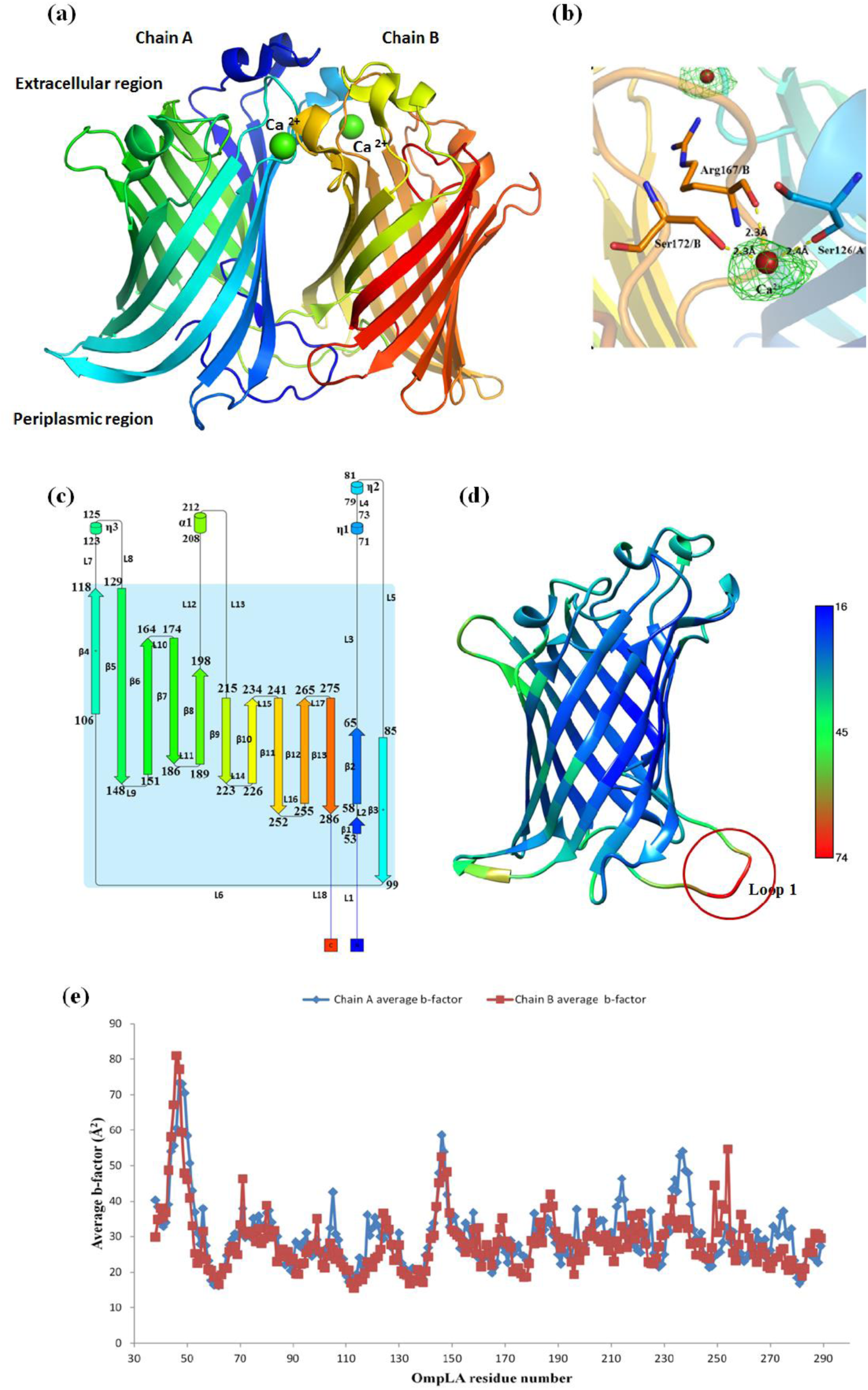
Crystal structure analysis of StOmpLA. **(a)** Three dimensional structure of calcium bound dimeric OmpLA, **(b)** Fo-Fc difference map (3δ) for two calcium ions along with coordination distances of interacting residues S172, R167 and S126, **(c)** Topology of OmpLA along with distribution and placement of 13 β-strands, 4 α-helices and 18 loops, **(d)** Average residue-wise temperature factor (Debye-Waller factor). Loop 1 shows highest temperature factor compare to rest of the structure **(e)** Comparison of temperature factors among chain A and B.

**Figure 2.**
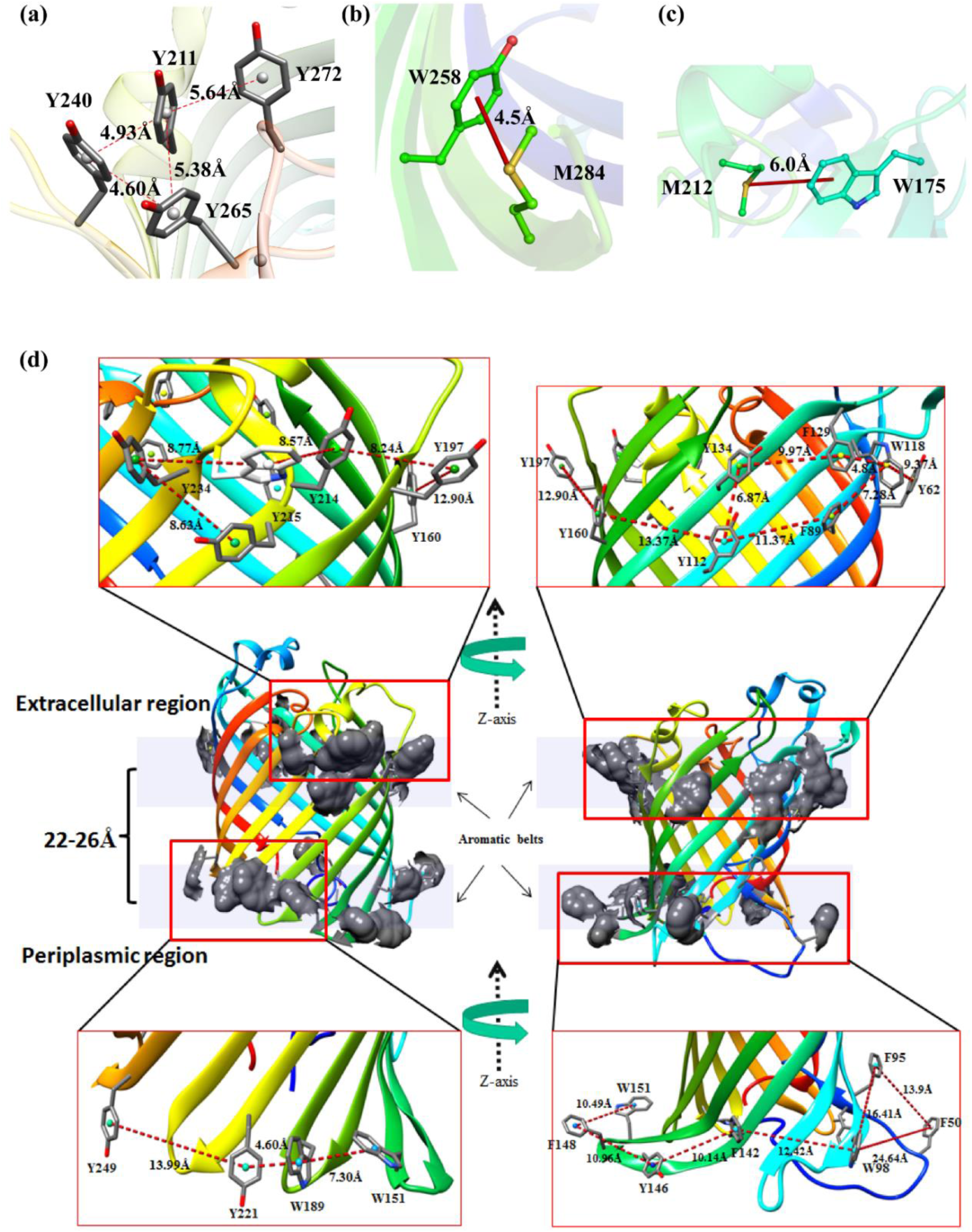
Structural features of StOmpLA involving aromatic amino acids. **(a)** Highly ordered aromatic ring cluster at extracellular end of the β-barrel involving Y211, Y240, Y265 and Y272, **(b)** and **(c)** show two sulphur-π interaction pairs between M284 and W258, and M212 and W175, respectively. **(d)** Two aromatic belts in OmpLA showing the arrangement of Tyr, Phe and Trp residues along periphery of β-barrel and distances between them are shown.

Crystal structure of StOmpLA has clear electron density in the region covering Q44 to T51 in both monomers, whereas density is missing for the corresponding region of EcOmpLA structure (E25 to F30). Totally 20 water molecules were modelled. We modelled two Ca^2+^ ions and one β-OG (n-Octyl-β-D-Glucopyranoside), used in crystallization buffer, using difference Fourier (Fo-Fc) densities. There is presence of only one β-OG detergent molecule in chain B towards the extracellular side of OmpLA with strong electron density for head region only, and each chain has 4 glycerol molecules (Fig. S3). Chain A has a total of 8 water molecules with 2 water molecules inside chain A channel while chain B has a total of 10 water molecules with 4 inside chain B channel (Fig. S3). Whether these channels are involved in transport of any solute is not known at present. β-OG was used in the final purification step while glycerol was present in the refolding and final elution buffers.

StOmpLA homodimer is stabilized by two calcium bridges. The calcium coordination in these two bridges have octahedral geometry for calcium bound to Ser126(O) of chain A with Arg167(O) and Ser172(OG) of chain B (50% vacancy)^17^, and trigonal bipyramidal geometry for calcium bridge with Ser126(O) of chain B, Arg167(O) and Ser172(OG) of chain A (40% vacancy)^17^. The low resolution of the StOmpLA may be the reason for absence of density for coordinating water molecules. Fo-Fc difference map, contoured at 3σ, shows electron density for two calcium ions bound at the dimer interface (Fig. 1b). Calcium binding is known to induce and stabilize the functional dimer formation of OmpLA. However, in *E. coli* OmpLA (1QD6) the first calcium is bound by octahedral geometry through C/S152, C/R147, D/S106 along with three water molecules (no vacancy). The second calcium is coordinated by trigonal bipyramidal geometry through C/S106, D/R147, D/S152 along with D/H_2_O302 (20% vacancy)^15^.

### Aromatic belts, dimer interface and crystal packing of StOmpLA

Each OmpLA monomer contains two aromatic belts around the β-barrel, separated by a distance of 22 to 26 Å on either side of the membrane which is very close to the average bacterial outer membrane thickness (Fig. 2d). The aromatic amino acids help anchor into membrane and stabilize the protein^18,19^. Aromatic side chains in these belts are in two major conformations; side chains towards the inner side of aromatic belts, located in the hydrophobic environment of detergent solubilized protein, are oriented away from polar solute, along the membrane plane, while the aromatic rings, particularly tyrosine with hydroxyl groups, located at the detergent-polar solvent interface are oriented towards the polar lipid head-solvent interface^20^ as seen in Figure 2d. Aromatic π-π interactions are implicated in the stability and self-assembly processes in proteins^20^. Three aromatic π-π interactions are noted between F129 and W118 (4.8 Å), Y134 and Y112 (6.87 Å), Y221 and W189 (4.60 Å). There are 12 tyrosine, 4 tryptophan and 6 phenylalanine residues marking the aromatic belt with a predominance of tyrosine residues. These Tyrosine residues contribute to the stability of OmpLA embedded in the outer membrane.

The dimer has a buried surface of 1429 Å^2^ which occludes 31% of the total solvent accessible area (PDBePISA)^21^. The dimer interface is also stabilized by the presence of 13 hydrogen bonds, 5 aromatic ring interactions and three hydrophobic patches as shown in Figure 3. The following hydrogen bonds are noted between A/N77(OD1): B/S168(OG) (3.7Å), A/N77(ND2): B/S168(OG) (3.3Å), A/S168(OG): B/N77(OD1) (3.3Å), A/S168(OG): B/N77(ND2) (3.7Å), A/s126(OG): B/D169(OD1) (2.8Å), A/S126(O): B/S172(OG) (3.5Å), A/G166(O): B/F129(N) (3.1Å), A/F129(N): B/G166(O) (2.8Å), A/Q144(OE1): B/Q144(NE2) (3.3Å), A/Q144(NE2): B/Q144(OE1) (3.2Å), A/P96(O): B/H254(N) (3.3Å), A/L52(O): B/L52(N) (2.8Å) and A/L52(N): B/L52(O) (2.8Å) residues. The hydrogen bond distribution is skewed with 8 at the extracellular end, 2 at the centre and 3 at the periplasmic end of dimer. Of the 8 hydrogen bonds at the extracellular side of the β-barrel, 6 (A/N77(OD1): B/S168(OG), A/N77(ND2): B/S168(OG), A/S168(OG): B/N77(OD1), A/S168(OG): B/N77(ND2), A/s126(OG): B/D169(OD1) and A/S126(O): B/S172(OG)) are positioned in polar non-membrane-embedded region of the protein dimer. The aromatic interactions which stabilize the dimer include A/Y134: B/F129 (4.0Å), A/F129: B/Y134 (4.1Å), A/Y112: B/F89 (5.8Å), A/F89: B/ Y112 (6.0Å) and A/Y53: B/H254 (5.9Å). The first four pairs of aromatic ring interactions are located in the centre of the dimer interface whereas the last interaction is located at the periplasmic end of the protein molecule. Figure 3 insets show these interactions with distances shown between the aromatic ring centroids. Three hydrophobic patches are formed towards the extracellular end, centre and periplasmic end of the β–barrel dimer as shown by the insets on right side of Figure 3. The extracellular patch consists of E125(A), W78(A), R167(A), S172(A), P128(A), S126(A), E125(B), W78(B), R167(B), S172(B), P128(B) and S126(B). The hydrophobic patch in the middle of dimer interface contains Y134(A), Y134(B), L91(A), L91(B), Y112(A) and Y112(B) whereas the patch towards the periplasmic end has L97(A), F95(A), Y53(A), P54(A), T51(A), L285(A), V255(A), T252(A), V255(B), L91(B), T51(B), P54(B), L93(B), F95(B), P96(B) and L97(B).

**Figure 3.**
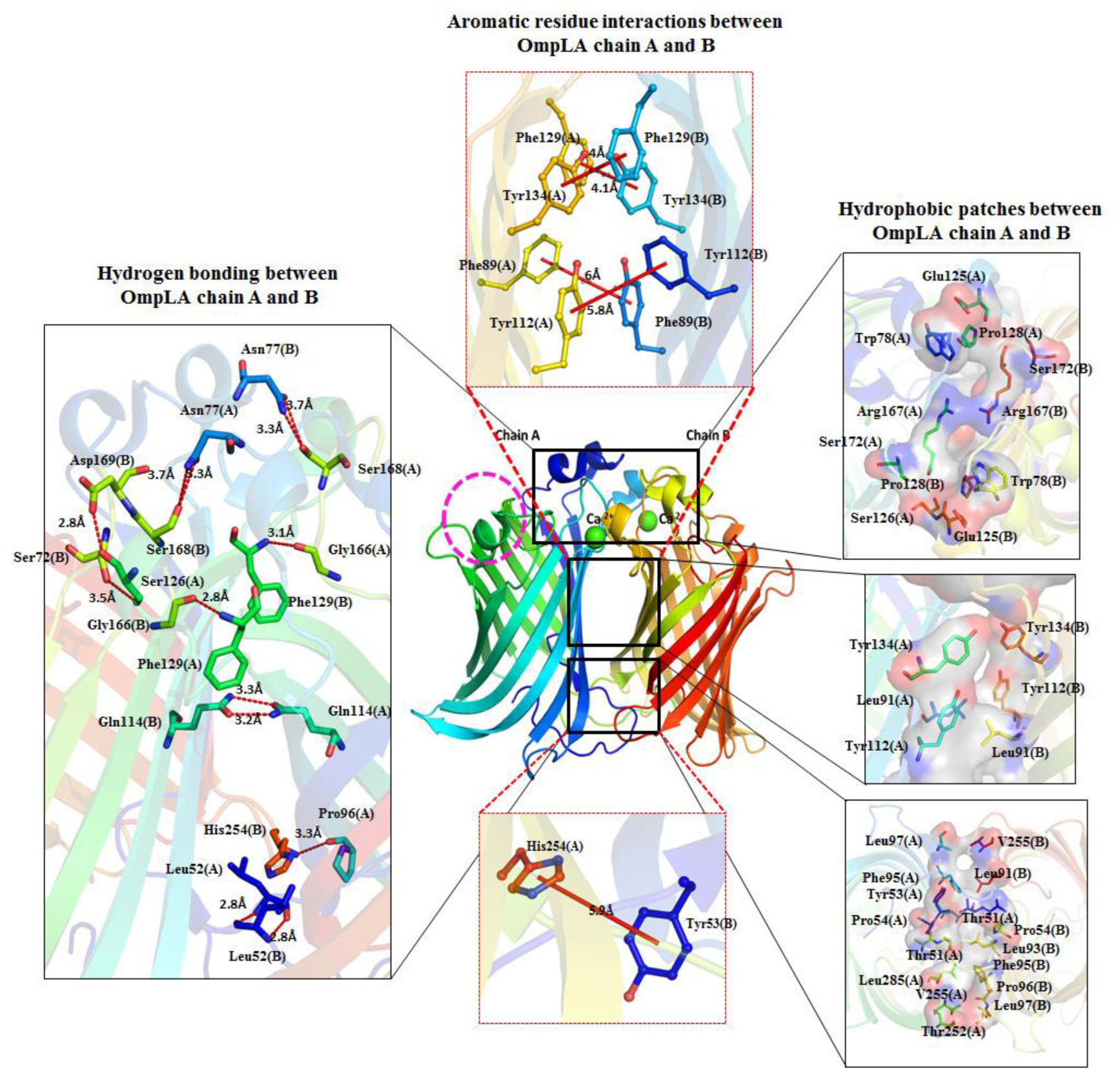
Structural analysis of OmpLA dimer interface. Left panel shows hydrogen bonding between two OmpLA chains, central panel shows aromatic ring interactions and right insets show the residues forming hydrophobic patches between two chains.

Crystals of OmpLA show typical type II packing^22^ wherein the detergent molecules shield hydrophobic transmembrane regions allowing crystal contacts to form through polar extra-membranous regions including loops and helices on both sides. The crystal contacts in OmpLA are shown in Figure S4 where the residues involved in crystal contacts are shown in all three planes. XY plane shows the alternate stacking orientation of OmpLA dimers in crystal. YZ plane clearly show the two regions of contact involving both highly convex sides of the protein. Two hydrophobic patches are formed by V281(A), V251(B’), Y62(A) and L257(B’) (Region I) as well as N176(B’), L178(B’), M158(A) and G103(A) (Region II). Region II also has a hydrogen bond between S201(B’) and L102(A) (3.22 Å). XZ plane also shows the presence of two more regions which help in crystal formation. Region III is formed by L70(A), E71(A), D67(A), N275(A), Y265(A), Y240(A), P206(B’’’), K210(B’’’) and N237(B’’’) while region IV mainly involves hydrophobic residues F148(B), A149(A), R147(B), L223(A’’) and G224(A’’). Region IV also has a hydrogen bond between E225(OE)(A’’) and amide backbone between F148(B) and A149(A) (2.9Å).

### Comparative analysis of *S. typhi* and *E. coli* OmpLA crystal structures

Structural comparison of StOmpLA monomer using DALI server showed best match with monomeric EcOmpLA structure (1QD5) with a z-score 39.8 and RMSD of 0.6 Å, while the dimeric structure of EcOmpLA 1QD6 shows RMSD of 0.36 Å. Both OmpLA proteins share 92% sequence identity, and the most variations are seen in the loops exposed to the extracellular space, turns facing cytoplasm and in all three 3_10_ helices (Fig. 4a b). Thus, the overall barrel topology and architecture of OmpLA from other Gram-negative human pathogens is expected to be conserved (Fig. 4b) including that of *S. flexneri*, mutation of which severely compromises type III secretion. The segment that comprises of residues 17 to 24 and 248 to 252 is α-helical in EcOmpLA but is as a loop in *S. typhi* OmpLA (Fig. 4a). Likewise, the segment comprising amino acid residues 70-74 in EcOmpLA is a loop but the corresponding segment is α-helical in *S. typhi* OmpLA though the sequences and positions are strictly conserved. Comparison of the monomeric and dimeric forms points to two very interesting changes: 1) monomeric OmpLA has two β-strands instead of one continuous β8-strand in *S. typhi* OmpLA and 2) the end of the β-strand has a very high b-factor asparagine (N181, 92.7 Å^2^) residue in 1QD5. The average b-factor for the loop and helix between β8 and β9 is also higher in monomeric forms in comparison to dimeric form of OmpLA. Moreover, the b-factor also varies with monomeric form having higher overall temperature factor when compared to dimeric form and distinctly higher B-factor in the extracellular helical regions (Fig. 4a). Also, there is a gradation in B-factor from 1QD5 > 1QD6 > 5DQX. The *S. typhi* OmpLA (5DQX – this study) shows a very low b-factor while as the EcOmpLA show higher average temperature factor.

**Figure 4.**
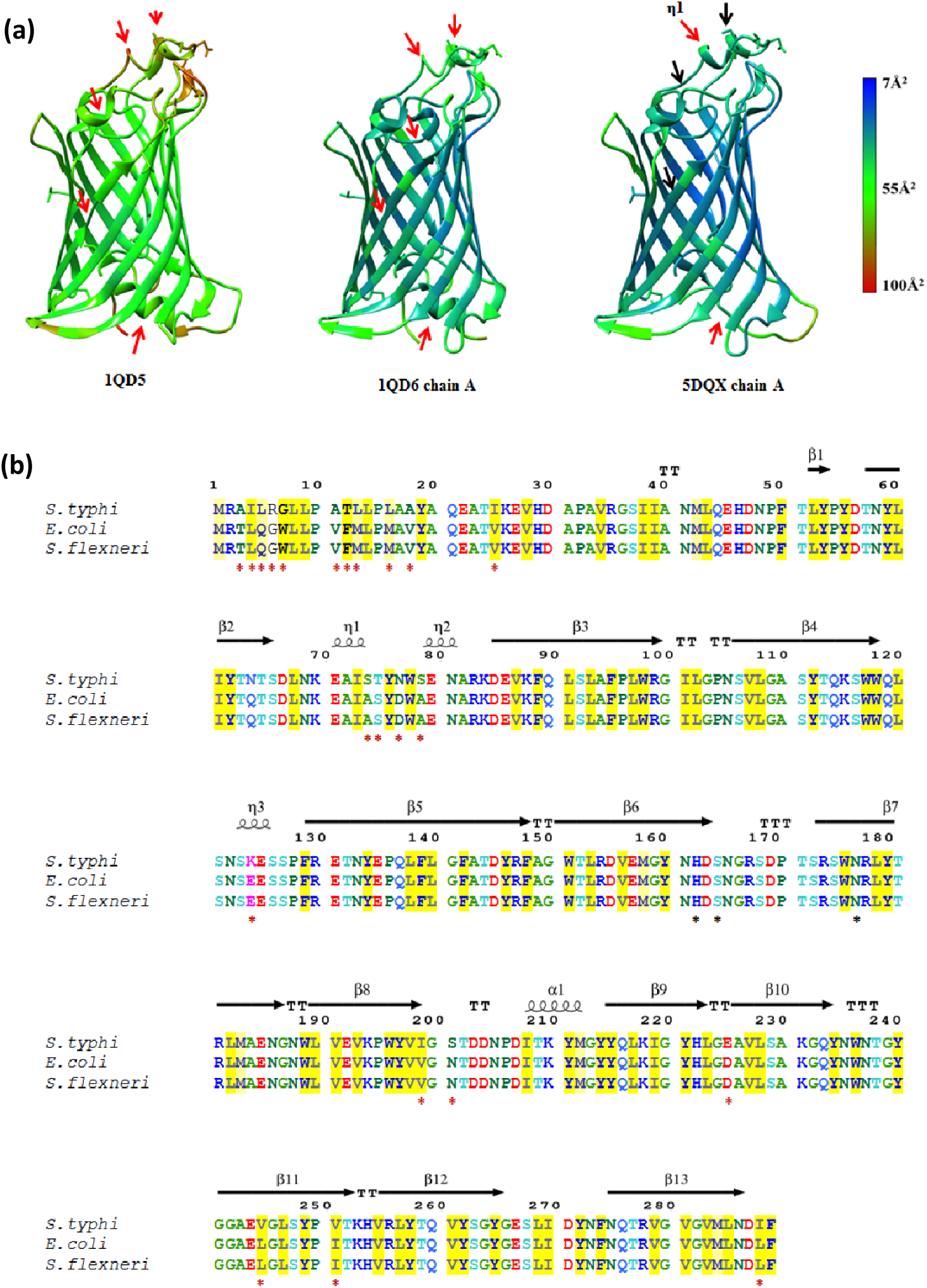
Comparison of crystal structures of OmpLA from *E. coli* (PDB:1QD5, 1QD6) and *S. typhi* (PDB:5DQX). **(a)** Temperature factor variation among monomeric and dimeric forms of OmpLA, **(b)** Structure based sequence alignment of OmpLA from *S. typhi, E. coli* and *S. flexneri*. Alignment is coloured based on 85% consensus using the following scheme: hydrophobic (ACFILMVWY), aliphatic (ILV) and aromatic (FHWY) residues shaded yellow; polar residues (CDEHKNQRST) are shaded blue; small (ACDGNPSTV) and tiny (AGS) residues shaded green; and big (QRKEILMWYF) residues shaded grey. OH group (ST) containing residues are shaded orange. Variations at the amino acid residue level are marked by red asterisk below them, and proposed active site residues of StOmpLA (162, 164 and 176) are marked by black asterisk. Red and black arrows indicate regions of higher and lower B-factors, respectively. Overall, StOmpLA has lower B-factors compare to monomeric and dimeric EcOmpLA.

### Calcium induced structural stability of StOmpLA

To assess the role of calcium in the stability and dynamics of StOmpLA, a 100 nanosecond molecular dynamics simulation (Desmond, Schrodinger suite) was performed with and without Ca^2+^. The RMSD plot shows that OmpLA with Ca^2+^ stabilizes faster than the protein without Ca^2+^, albeit at a higher RMSD value. RMSD trajectory for OmpLA without Ca^2+^ stabilize towards the end of 60 ns and overlaps the native OmpLA trajectory (Fig. 5a). Most of the RMSD fluctuations were seen in the loop regions as marked by the red boxes (Fig. 5b) with loop lengths having no bearing on the RMSF values as seen for loop regions L3, L4 and L5 (Fig. 1c). The Ca^2+^ binding residues Ser126/A along with Ser167/B and Arg172/B show higher RMSF compared to second calcium binding residues, Ser126/B, Ser167/A and Arg172/A, clearly visible in box III and VI. Higher RMSF of Ca^2+^ binding residues were observed in an earlier study as well^23^. The higher RMSF in Ca^2+^ minus state shown in boxes I and IV can be explained by absence of calcium coordinated water mediated hydrogen bonding network as well as weakening of hydrogen bonding involving 4 out of 6 hydrogen bonds on the extracellular side of β-barrel namely A/N77(OD1): B/S168(OG), A/N77(ND2): B/S168(OG), A/S168(OG): B/N77(OD1), A/S168(OG): B/N77(ND2). Overall, the dynamics analysis suggests that calcium bound dimer is stable compare to the unbound structure. The dimeric structure of StOmpLA with bound calcium was further used as a template for following *in silico* study.

**Figure 5.**
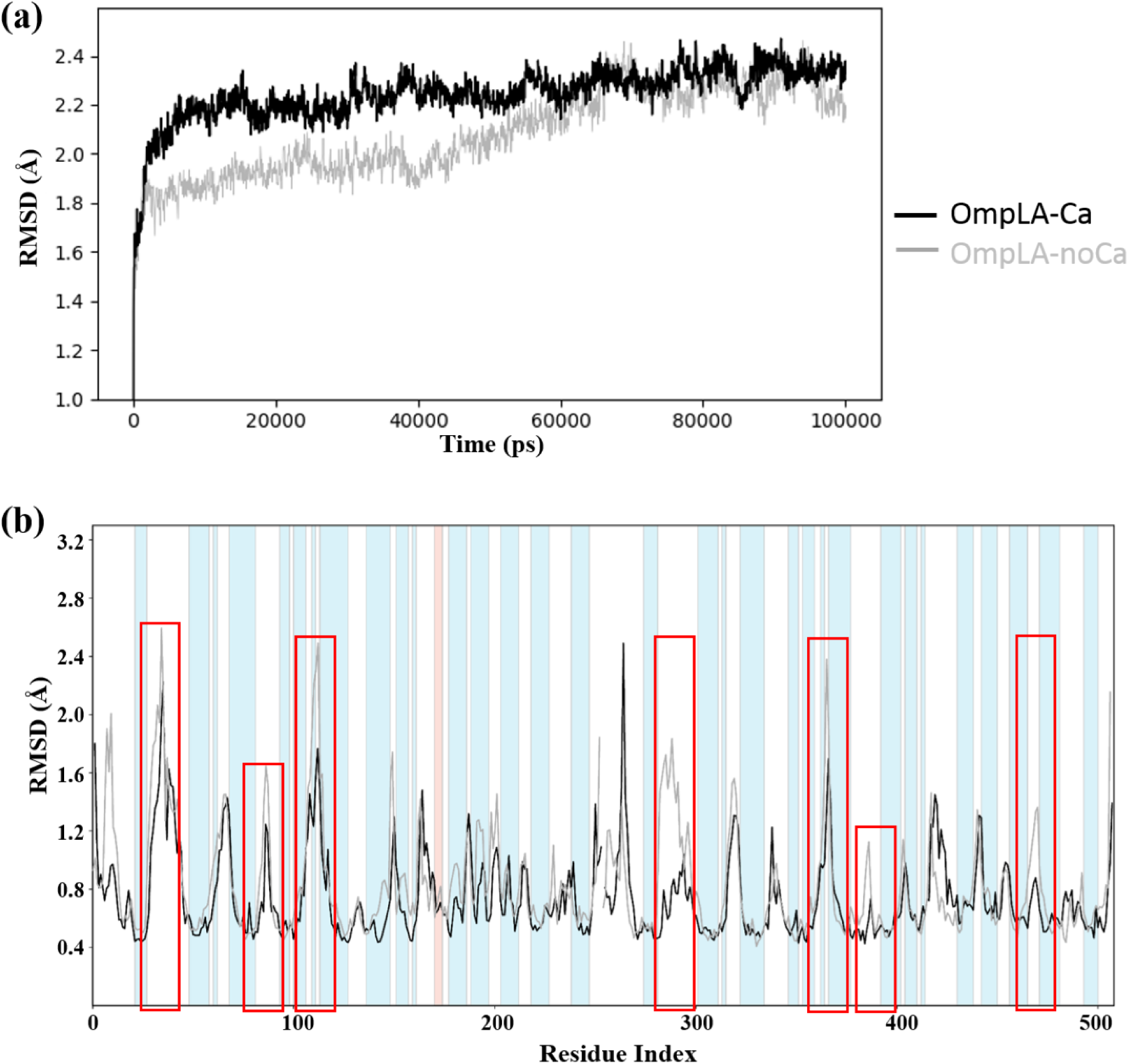
Molecular dynamics analysis StOmpLA. **(a)** Comparative structural stability analysis of OmpLA with and without Ca^2+^, black and grey lines respectively, subjected to 100 ns simulation and corresponding RMSF comparison **(b)**. Regions with high variations are marked by red boxes.

### *In silico* structure-based anti-microbial discovery targeting *S*tOmpLA

To target StOmpLA with small molecular inhibitors, a thorough structural analysis of binding pockets was done using the SiteMap module of Schrodinger suite, which predicts druggable pockets, based on size, shape and chemical features (Table S1). The top ranked site (site-1) is found at the dimer interface, facing the extracellular side of (Fig. 6a, b), and contains the following residues from both chains A & B: 75-78, 128-132,134, 165 −171. Site-1 is exposed to solvent from the extracellular side of bacterial outer membrane. Thus, further *in silico* screening was carried out targeting site-1 using a set of synthetic compounds, phytochemicals, NCI and FDA approved drug database compounds. Potent binders were identified and short-listed based on the G-score (Schrodinger: Glide) and number of hydrogen bonds and hydrophobic interactions within the predicted site. Compounds with Glide scores ranging from −13.2 to −9.8 are listed in Table S2. Site-2, located right beneath site-1 in each monomer, spans the buried interior space between the two monomers where the native membrane lipid substrate is expected to bind. This is evident from the complex crystal structure of EcOmpLA with covalently bound inhibitor hexadecanesulfonyl fluoride (HDSF)^24^. Binding of two molecules of HDSF at the largely hydrophobic dimer interface suggest that this site can accommodate two molecules of hydrophobic inhibitor targeted to bind at the buried dimer interface, though bioavailability and toxicity such molecule remains to be tested experimentally at the context of known HSDF toxicity. Site-3 and Site-4 are equivalent sites present inside the interior channel-like opening of each monomer, closer to the middle of the barrel height. Site-5 spans the both monomers from the periplasmic space side and is considered less druggable (not shown).

**Figure 6.**
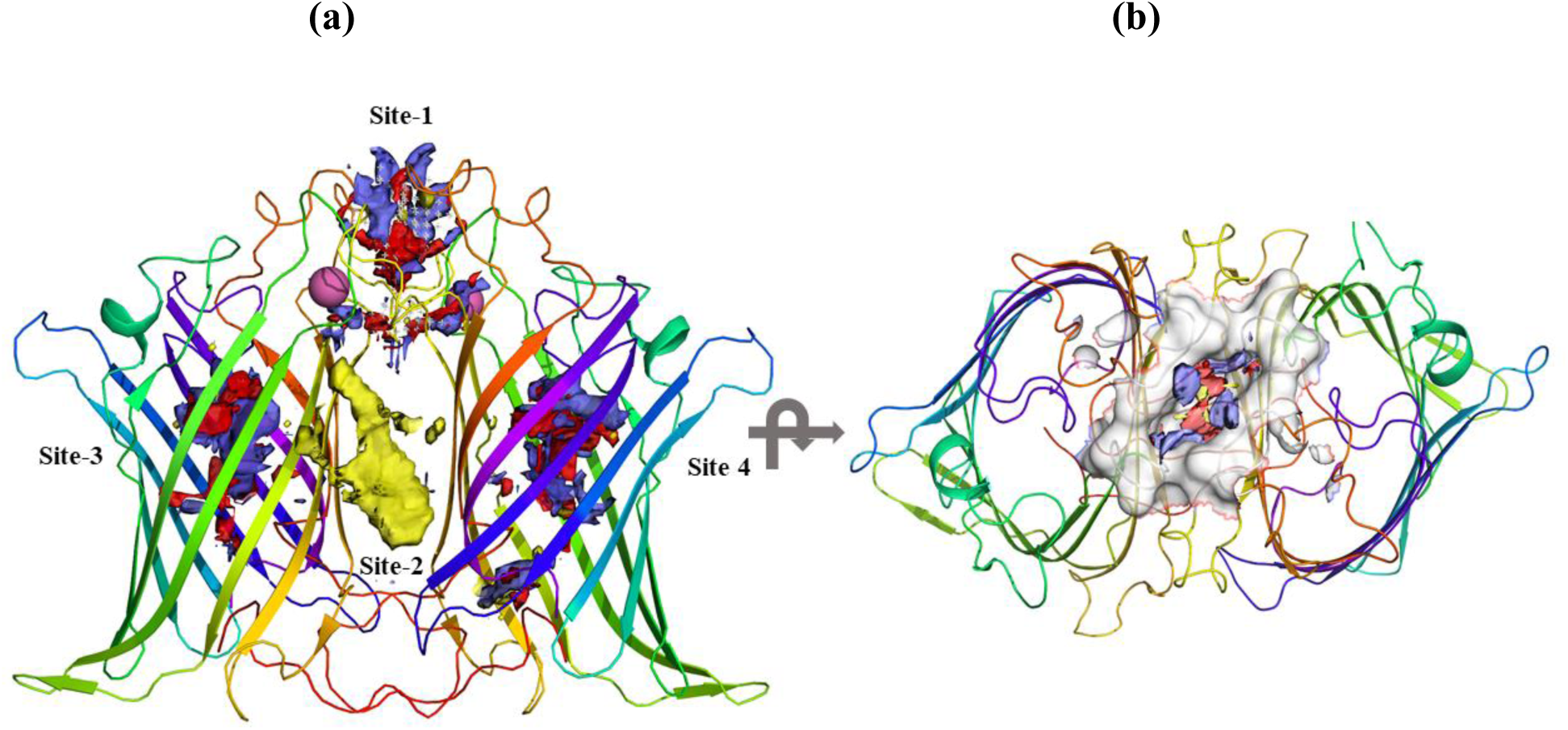
Druggable binding pockets in stOmpLA predicted using SiteMap (Schrodinger). **(a)** Out of the five sites predicted, only four are shown here. Site-2, made of mostly flat and hydrophobic, has an equivalent site on opposite side of the dimer, not shown here, **(b)** Site-1 seen from the extracellular space, towards periplasmic side. Druggable pocket characteristics are color coded differently: Hydrogen donor; blue, Hydrogen acceptor; red, Hydrophobic; yellow.

The top ranked *in silico* hits binding to site-1; NCI97317, Alanylthreonine and Phloretin were further explored for structural stability using molecular dynamics using 100ns simulations. RMSD values of the protein-ligand systems were compared with reference to the initial protein structure. The RMSD time course trajectories for four complexes are shown with native OmpLA dimer as the control in Figure 7. The initial fluctuating RMSD trajectories approached stable values towards the end of 100 ns MD simulations, indicating equilibrated protein-ligand complexes, suitable for various analysis. The RMSD values were found to be lower in OmpLA-small molecule complexes. Variations of RMSD, in comparison to native protein, along with representative hydrogen bonding pattern in the stable trajectory region, as insets, are shown in Figure 7. OmpLA-NCI97317 complex showed the least variation in RMSD among the four complexes analysed. The complex is stabilized by three water mediated hydrogen bonds from A/F129, B/W78 and B/F129 along with two hydrogen bonds with A/W78 (Fig. 7). OmpLA-alanylthreonine complex is stabilized by 5 hydrogen bonds from B/N77, B/T75 and B/Y76 whereas OmpLA-phloretin complex is stabilized by 4 hydrogen bonds with A/Y134, A/N165, A/R167 and B/Y76. OmpLA-sulphamethoxaole complex is stabilized by 5 hydrogen bonds with A/R167, BY76, B/E131 and B/N 165 along with one water mediated hydrogen bond with A/F129. Among all the complexes, the RMSD values for OmpLA-alanylthreonine complex has higher fluctuation values ranging between 1.12 and 2.0 Å. The other three inhibitor complexes have RMSD values varying between 1.4 and 1.8 Å. The RMSF values, for all the protein-inhibitor complexes, stabilize for the extracellular part of barrel covering region between loops L3 and L5 marked by red box (Fig. S5). OmpLA/inhibitor complex interactions, categorized into hydrogen bonds, ionic, hydrophobic and water bridges, and monitored throughout the 100ns simulation are shown in Figure S6. Residues with values more than 1 make multiple contacts with these potential binders. A detailed 2D representation of an elaborate interaction pattern for more than 30%-time occupancy during the entire 100ns simulation is shown in Figure S7. These results clearly indicate the structural stability of docked complexes and further suggest that StOmpLA is druggable. Further experimental validation, using OmpLA enzyme inhibition assay *in vitro*^24^, will help design unique inhibitors of StOmpLA.

**Figure 7.**
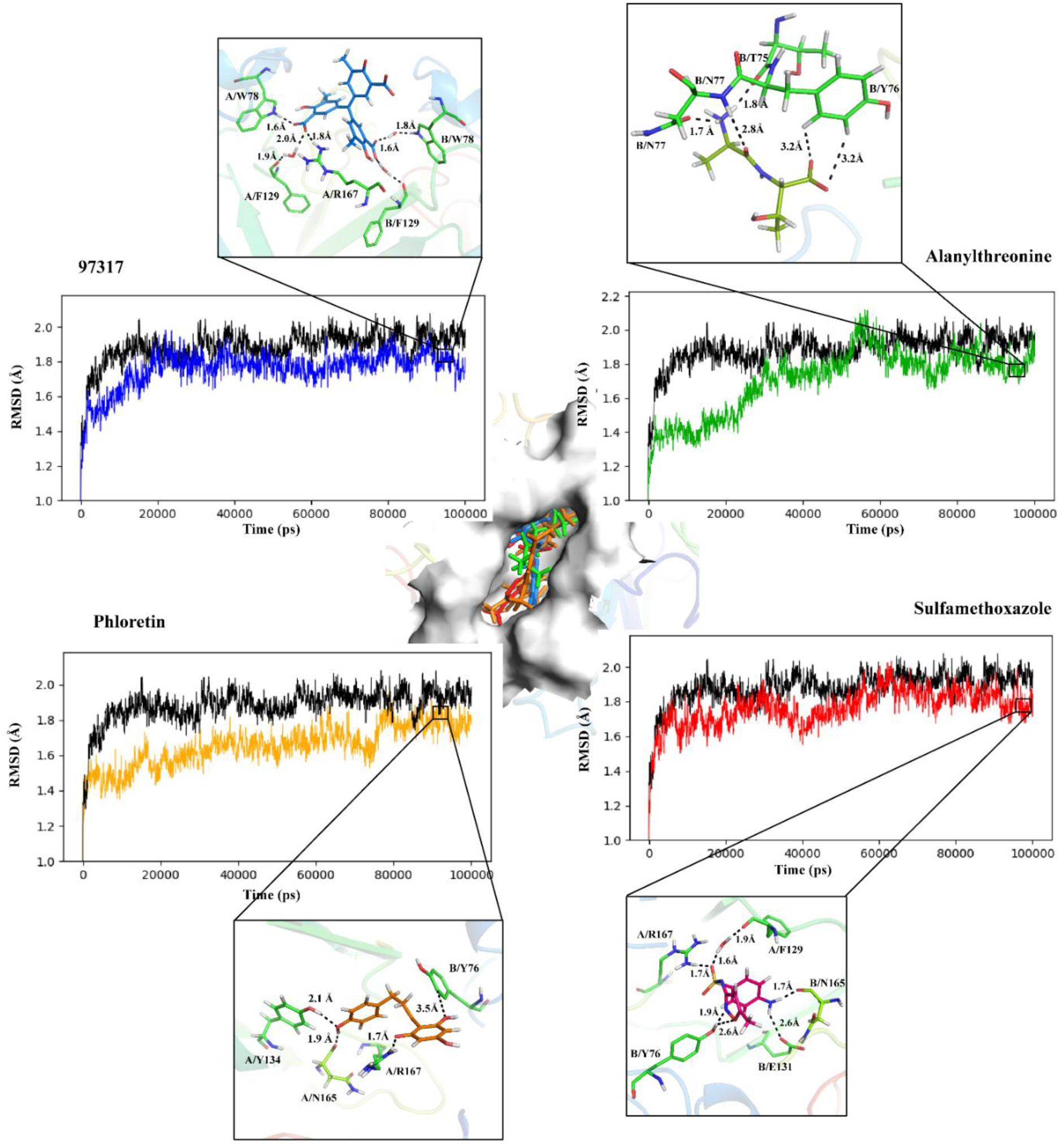
Molecular dynamics simulation analysis of OmpLA-inhibitor complexes docked *in silico*. Docking of top three hits and a known antibiotic Sulfamethoxazole at the site-1 are shown at the centre. RMSD trajectories for all four complexes, in comparison to native protein are shown. Representative hydrogen bonding pattern is shown, as insets, for each OmpLA-small molecule complex for most stable trajectory region on the RMSD plots.

## Summary

The crystal structure of calcium bound StOmpLA was determined to the resolution 2.95 Å. The functional dimeric structure was used as a template to screen potential small molecule binders that target the top ranked druggable pocket in the dimer interface of StOmpLA. Docked complexes of top three hits; NCI97317, Alanylthreonine and Phloretin from 14 shortlisted compounds were assessed for structural stability using 100 ns molecular dynamics simulations. The data presented here provides a framework for further experimental validation that will help develop therapeutics specifically targeting virulence causing mechanism of Gram -negative pathogens, encoding OmpLA. This approach may help address the growing problem of antibiotic resistance.

## Methods

### Cloning of StOmpLA encoding *pldA* gene

810 bp long *pldA* gene, encoding leaderless StOmpLA (21Q-289F) was PCR amplified using *S. typhi* Ty21a genomic DNA as a template, with an annealing temperature gradient of 48°C to 52°C. PCR primers used were; forward-5’GCCATATGCAAGAAGCTACGATAAAAG 3’, reverse-5’GCGGATCCTCAGAAGATATCGTTAAG3’. Maximum amplification was observed at 50°C annealing temperature (Fig. S1a) and cloned between *Nde*I and *Bam*HI restriction sites into the pET-30b vector, after restriction digestion and ligation. Ligation mixer was used to transform DH5α cells and positive clones were identified by colony PCR, confirmed by restriction digestion with *Nde*I and *BamH*I enzymes (Fig. S1b), and followed by DNA sequencing.

### Overexpression, refolding and purification of StOmpLA

*E. coli* T7 Express/I^q^ cells were used for overexpression. Single colony was inoculated and grown overnight at 37°C in Luria Bertani (LB) (Himedia Labs), supplemented with 30µg/ml kanamycin. 1% of the overnight grown cells were subcultured and induced at 37°C for 16 hours in 1 L of LB-AIM (LB Auto Induction Media). Cells were harvested by centrifugation at 4500g for 20 min at 4°C and stored at −20°C. The signal peptideless StOmpLA was seen in the inclusion bodies (IBs), similar to other overexpressed outer membrane proteins without signal peptide. Cell pellet was resuspended in 50 mM Tris HCl (pH 8.0) and sonicated at 80Hz for 20 minutes with cycles of 3 seconds ON and 9 seconds OFF. Cell lysate was centrifuged at 10000 rpm (rotor # 3335, Heraus) for 7 min at 20°C to collect inclusion bodies (IBs), unlysed cells and cell debris. IBs were washed three times with a buffer containing 25 mM Tris HCl (pH 8.3), 0.1 M NaCl and 2% Triton X-100, and 2 M urea, followed by two washes using the buffer containing 25 mM Tris-Cl (pH 8.3) and 0.1 M NaCl. At each step, IBs were resuspended using Dounce homogenizer and then kept on rotary shaker at 37°C for 15 minutes followed by centrifugation at 7,000 rpm (rotor # 3335, Heraus) for 7 minutes at 20°C. Typical yield of purified inclusion bodies were 1 gram per litre of culture.

### Unfolding, refolding and purification of StOmpLA

Purified IBs were solubilized in Tris-HCl buffer containing varying concentrations of Urea to choose the final concentration for unfolding. Final, large-scale unfolding was carried out in the buffer containing 25 mM Tris HCl (pH 8.3), 0.1 M NaCl and 8 M urea for 3 hours at 37°C, with moderate shaking^25^. Unfolded OmpLA was centrifuged at 13,000 rpm (rotor # 3335, Heraus) for 45 minutes at 25°C followed by passing through 0.45 m filter to remove particulate matters. Refolding was done by slow (drop by drop) dilution into 10-fold volume of refolding buffer containing 25 mM Tris HCl (pH 8.3), 0.1 M NaCl, 10% (v/v) glycerol and 0.3% C_12_E_9_ (Sigma), at a flow rate of ∼ 25ml/h at 20°C for 16 hours with moderate stirring to ensure maximum refolding^25^. The diluted and refolded protein was concentrated to 20 ml using ultrafiltration Amicon stirred cell (Millipore) attached with a 10 kDa MWCO membrane (Stirred cell and Centriprep-10), and centrifuged at 13,000 rpm (rotor #,FA-45-30-11, Eppendorf) for 45 minutes at 20°C to remove small aggregates and particulate matter.

Refolded OmpLA was diluted 10-fold into a buffer containing 25 mM Tris HCl (pH 8.3) and 0.3% C_12_E_9,_ and loaded onto a 5ml HiTrap Q-HP anion-exchange column (GE). Column equilibration and washing, after sample loading, was done using 25 mM Tris HCl (pH 8.3), 10 mM NaCl and 0.3% C_12_E_9_. Bound protein was eluted, in steps, with the same buffer containing 1 M NaCl and checked on denaturing and reducing SDS-PAGE. Pooled samples after Q-HP column was concentrated and loaded onto a preparative Superdex 200 10/300 column, attached to an AKTA Explorer (GE), pre-equilibrated with 25 mM Tris HCl (pH 8.3), 0.01 M NaCl and 0.3 % (v/v) C_12_E_9_. The fractions containing pure OmpLA were pooled and further passed through a 1ml HiTrap Q-Sepharose fast flow (GE healthcare) anion-exchange column equilibrated with the same buffer, at the flow rate of 0.3ml/min. Column was washed with the wash buffer containing 25 mM Tris HCl (pH 8.3), 10 mM NaCl, and 1% β-OG, on a loop, for 3 hours at 4°C to remove C_12_E_9_ and unbound protein. Then, the final round of column wash done with wash buffer containing 1 % β-OG for overnight. After detergent exchanged protein was eluted in a single step using Buffer-B containing 25 mM Tris HCl pH 8.3, 1 M NaCl, 10% glycerol, 1% β-OG. Eluted protein was concentrated and salt concentration was reduced using Amicon centriprep-10 ultracentifugal devise. Aliquots of 40 µl of purified protein were flash frozen in liquid nitrogen and stored at −80 °C until further use.

### Circular dichroism (CD)

CD measurements were performed on a Jasco J-810 spectropolarimeter. For far-UV CD spectra of secondary structure, samples in 1 mm path length cuvettes were scanned in the wavelength range 250–190 nm, using a 1 nm nominal bandwidth with three accumulations. Spectra were corrected for background by subtraction of buffer and detergent blanks.

### Crystallization

OmpLA crystallization trials were carried out using sitting drop with the nanovolume dispensing robot (Mosquito, TTP Labtech Ltd) and various commercial sparse matrix crystal screens like JCSG (+), PACT Premier (Molecular Dimensions), Wizard I/II (Emerald Biosystems), Crystal Screen I/II (Hampton Research), MemGold and MemStart. 300 nl drops were set up with three (1:1, 1:2, 2:1) ratios of protein to crystallizing buffer and incubated at 293 K/20 °C. A protein concentration of 14mg/ml was used for crystallization. Needle shaped crystals were obtained after a day in a condition containing 0.2 M calcium chloride, 0.1 M sodium acetate (pH 5.0), 20% w/v PEG 6000 whereas micro-crystals appeared after a day in a different condition; 0.08 M sodium citrate pH 5.2, 2.2 M ammonium sulphate and 0.64 M sodium acetate, 18% w/v PEG3350. Crystals were observed under a light microscope (Olympus). Diffraction quality 3D crystals grew within two days in 0.1 M sodium iodide, 0.1 M sodium phosphate (pH 7.0), and 33% v/v polyethylene glycol 300.

### X-ray diffraction data collection

The crystal was swiftly fished out from the mother liquor using a nylon-fibre loop after adding 2 µl of reservoir solution containing 0.1 M sodium iodide, 0.1 M sodium phosphate (pH7.0), and 33 % v/v polyethylene glycol 300. X-ray diffraction data was collected to 2.95Å resolution on an in-house rotating anode X-ray source (Rigaku FR-E+ Super Bright) connected to R-AXIS IV++ detector at the National Institute of Immunology (NII), New Delhi, India. A total of 155 images were collected at the wavelength of 1.5418 Å with 2 min exposure and 1° oscillation per image at 100K.

### Structure determination, refinement and analysis

Data was indexed, integrated and scaled using the HKL-2000 software package^26^ and automated Molecular Replacement was performed using BALBES server^27^. Initial rigid body and restraint refinements were performed using REFMAC5^28^ of CCP4 suite^29^ and model was built using WinCoot0.7.2.1^16^. The final protein model was validated using PROCHECK module of CCP4 suite. Structure-based multiple sequence alignment was done using the ESPript server^30^. PyMOL was used for structure visualization, comparison and generating figures.

### Compounds retrieval from publicly available databases

Compounds with known anti-biotic activity; Ampicillin, Chloramphenicol, hexadecanesulfonyl fluoride, Kanamycin, ONO-RS-082, Streptomycin, Daptomycin, 8-Methoxy Fluoroquinolone, aristolochic acid I, Azithromycin, bromoenol lactone, Ciprofloxacin, Clarithromycin, Clindamycin, Clofazimine, Dapsone, erythromycin, Ethambutal, Gatifloxacin, Gentamycin, Halopemide, Levofloxacin, Linezolid, Methyl linolenyl fluorophosphonate, Moxifloxacin, Neomycin, Nitrofuranton, Nortoxacin, Rifabutin, Rifapentine, Sulfacetamide, telavacin, tigecycline, Trimethoprim, sulfamethoazole, Norfloxacin, Isoniazid and phytochemicals; Allicin, Artemisin, Asiatocoside, Berberine, Caffeic acid, Capsaicin, Catechin, Chrysin, Cocaine, Coumarin, Ellagitannin, Eugenol, Fructose, harmane, p-benzoquinone, Phloretin, Protoanemonin, Salicylic acid, Terebinthone, Tobramycin, and Withafarin from PubChem database (http://www.ncbi.nlm.nih.gov/pccompound), 3,11,428 compounds from NCI database (https://cactus.nci.nih.gov/download/nci/index.html) and 61,178 compounds from FDA Approved Drug database were used for *in silico* screening studies.

### Molecular Docking

Crystal structure of StOmpLA was prepared using “protein preparation” wizard of Schrodinger suite version 2018-3, Licensed to ICGEB, New Delhi, to relieve steric clashes using the OPLS3e force field^31^. Small molecules were prepared by LigPrep module to expand protonation and tautomeric states at 7.0±2.0 pH. Grid was generated for the site-1 predicted and scored by SiteMap. Molecular docking was carried out using Glide.

### MD simulations

System Builder module from Schrodinger’s Maestro was used to set up a POPC membrane system at 300K. All the systems were constructed using Desmond MD package using OPLS3e force field to calculate the atomic interactions^31^. An orthorhombic box with a box volume of 2111363 Å^3^ with buffer distances of 10 Å on each vertex was used to submerge the protein or protein-inhibitor complexes. The default simple point charge (SPC) water model was used with 0.15 M NaCl to neutralize the system. Sequentially, 2000 iterations of minimization were performed to bring the system into local energy minima with a convergence threshold of 1 kcal/mol/Å using Desmond. The cut-off radius for the short-range Coulombic interactions was set to 9 Å. Molecular dynamics simulation was performed on the relaxed model system at 300 K and 1.01325 bar pressure for 100 ns using NPT ensemble with the recording interval of 50 ps for trajectory and 1.2 ps for energy by Nose-hoover thermostat method. Post-processing was done by using Simulation Quality Analysis, Simulation Event Analysis and Simulation Interaction Diagram tools to analyse the protein stability using RMSD, RMSF, hydrogen bonding, hydrophobic interactions, π-π stacking, salt-bridge interactions and energy parameters.

## Acknowledgement

The authors thank Muthusankar Aathi, DST-NPDF, for assistance with *in silico* screening, S. Krishnaswamy and D. Balasubramnian for critical reading of the manuscript, Bichitrakumar Biswal, Paul Ravikant, NII, New Delhi for help with X-ray data collection, Amit Kumar and Manojkumar, Madurai Kamaraj University, for assistance in data collection and processing, respectively, and Vinod Devaraji, Schrodinger-India for help with MD simulations. SBN lab is funded by ICMR and Bharathiar University. PP and RR were supported by fellowships from DST-NPDF (PDF/2016/003347) and CSIR, respectively. Schrodinger suite is supported by ICGEB core funds. Research in AA lab is funded by ICGEB core funds and grants from Department of Biotechnology, Govt. of India; BT/PR13735/BRB/10/786/2011 and BT/PR28080/BID/7/836/2018.

## Author contribution

NSB and AA conceived and supervised the work, PP cloned, refolded, purified, and crystallized StOmpLA. PP, RR and AA collected X-ray data and determined the structure, PP and RR performed *in silico* studies. All the authors contributed to writing the manuscript.

## References

1 Homma, H. et al. The DNA sequence encoding pldA gene, the structural gene for detergent-resistant phospholipase A of E. coli. Journal of biochemistry 96, 1655–1664 (1984).

2 Brok, R. G. et al. Molecular characterization of enterobacterial pldA genes encoding outer membrane phospholipase A. Journal of bacteriology 176, 861–870 (1994).

3 Horrevoets, A. J., Hackeng, T. M., Verheij, H. M., Dijkman, R. & de Haas, G. H. Kinetic characterization of Escherichia coli outer membrane phospholipase A using mixed detergent-lipid micelles. Biochemistry 28, 1139–1147 (1989).

4 Pugsley, A. P. & Schwartz, M. Colicin E2 release: lysis, leakage or secretion? Possible role of a phospholipase. The EMBO journal 3, 2393–2397 (1984).

5 Pullen, J. K., Liang, S. M., Blake, M. S., Mates, S. & Tai, J. Y. Production of Haemophilus influenzae type-b porin in Escherichia coli and its folding into the trimeric form. Gene 152, 85–88 (1995).

6 Audet, A., Nantel, G. & Proulx, P. Phospholipase A activity in growing Escherichia coli cells. Biochimica et biophysica acta 348, 334–343 (1974).

7 Cronan, J. E., Jr. & Wulff, D. L. A role for phospholipid hydrolysis in the lysis of Escherichia coli infected with bacteriophage T4. Virology 38, 241–246 (1969).

8 de Geus, P., van Die, I., Bergmans, H., Tommassen, J. & de Haas, G. Molecular cloning of pldA, the structural gene for outer membrane phospholipase of E. coli K12. Molecular & general genetics : MGG 190, 150–155 (1983).

9 Wang, X. et al. The outer membrane phospholipase A is essential for membrane integrity and type III secretion in Shigella flexneri. Open biology 6, 160073 (2016).

10 Schmiel, D. H. & Miller, V. L. Bacterial phospholipases and pathogenesis. Microbes and infection 1, 1103–1112 (1999).

11 Rosenbusch, J. P. Characterization of the major envelope protein from Escherichia coli. Regular arrangement on the peptidoglycan and unusual dodecyl sulfate binding. The Journal of biological chemistry 249, 8019–8029 (1974).

12 Tokunaga, M., Tokunaga, H., Okajima, Y. & Nakae, T. Characterization of porins from the outer membrane of Salmonella typhimurium. 2. Physical properties of the functional oligomeric aggregates. European journal of biochemistry 95, 441–448 (1979).

13 Park, K., Perczel, A. & Fasman, G. D. Differentiation between transmembrane helices and peripheral helices by the deconvolution of circular dichroism spectra of membrane proteins. Protein science : a publication of the Protein Society 1, 1032–1049, doi:10.1002/pro.5560010809 (1992).

14 Snijder, H. J. et al. Structural evidence for dimerization-regulated activation of an integral membrane phospholipase. Nature 401, 717–721, doi:10.1038/44890 (1999).

15 Snijder, H. J. et al. Structural investigations of calcium binding and its role in activity and activation of outer membrane phospholipase A from Escherichia coli. Journal of molecular biology 309, 477–489, doi:10.1006/jmbi.2001.4675 (2001).

16 Emsley, P. & Cowtan, K. Coot: model-building tools for molecular graphics. Acta crystallographica. Section D, Biological crystallography 60, 2126–2132, doi:10.1107/S0907444904019158 (2004).

17 Zheng, H. et al. Validation of metal-binding sites in macromolecular structures with the CheckMyMetal web server. Nature protocols 9, 156–170, doi:10.1038/nprot.2013.172 (2014).

18 Seshadri, K., Garemyr, R., Wallin, E., von Heijne, G. & Elofsson, A. Architecture of beta-barrel membrane proteins: analysis of trimeric porins. Protein science : a publication of the Protein Society 7, 2026–2032, doi:10.1002/pro.5560070919 (1998).

19 Ulmschneider, M. B. & Sansom, M. S. Amino acid distributions in integral membrane protein structures. Biochimica et biophysica acta 1512, 1–14 (2001).

20 Madhusudan Makwana, K. & Mahalakshmi, R. Implications of aromatic-aromatic interactions: From protein structures to peptide models. Protein science : a publication of the Protein Society 24, 1920–1933, doi:10.1002/pro.2814 (2015).

21 Krissinel, E. & Henrick, K. Inference of macromolecular assemblies from crystalline state. Journal of molecular biology 372, 774–797, doi:10.1016/j.jmb.2007.05.022 (2007).

22 Russo Krauss, I., Merlino, A., Vergara, A. & Sica, F. An overview of biological macromolecule crystallization. International journal of molecular sciences 14, 11643–11691, doi:10.3390/ijms140611643 (2013).

23 Baaden, M., Meier, C. & Sansom, M. S. A molecular dynamics investigation of mono and dimeric states of the outer membrane enzyme OMPLA. Journal of molecular biology 331, 177–189 (2003).

24 Horrevoets, A. J., Verheij, H. M. & de HAAS, G. H. Inactivation of Escherichia coli outer-membrane phospholipase A by the affinity label hexadecanesulfonyl fluoride: Evidence for an active-site serine. European journal of biochemistry 198, 247–253 (1991).

25 Balasubramaniam, D., Arockiasamy, A., Kumar, P. D., Sharma, A. & Krishnaswamy, S. Asymmetric pore occupancy in crystal structure of OmpF porin from Salmonella typhi. Journal of structural biology 178, 233–244, doi:10.1016/j.jsb.2012.04.005 (2012).

26 Otwinowski, Z. & Minor, W. Processing of X-ray diffraction data collected in oscillation mode. Methods in enzymology 276, 307–326 (1997).

27 Long, F., Vagin, A. A., Young, P. & Murshudov, G. N. BALBES: a molecular-replacement pipeline. Acta crystallographica. Section D, Biological crystallography 64, 125–132, doi:10.1107/S0907444907050172 (2008).

28 Murshudov, G. N. et al. REFMAC5 for the refinement of macromolecular crystal structures. Acta crystallographica. Section D, Biological crystallography 67, 355–367, doi:10.1107/S0907444911001314 (2011).

29 Winn, M. D. et al. Overview of the CCP4 suite and current developments. Acta crystallographica. Section D, Biological crystallography 67, 235–242, doi:10.1107/S0907444910045749 (2011).

30 Robert, X. & Gouet, P. Deciphering key features in protein structures with the new ENDscript server. Nucleic acids research 42, W320–324, doi:10.1093/nar/gku316 (2014).

31 Harder, E. et al. OPLS3: A Force Field Providing Broad Coverage of Drug-like Small Molecules and Proteins. Journal of chemical theory and computation 12, 281–296, doi:10.1021/acs.jctc.5b00864 (2016).

