## Supplementary table S1, figures S1- to S7 for "Crystal structure of calcium bound outer membrane phospholipase A (OmpLA) from *Salmonella typhi* and *in silico* anti-microbial screening"

**Communicating author**:

Arockiasamy Arulandu

Membrane Protein Biology Group, International Centre for Genetic Engineering and Biotechnology (ICGEB),

Aruna Asaf Ali Marg,

New Delhi 110067. India.

**Table S1**: Prediction and ranking of druggable pockets in StOmpLA dimer using SiteMap

| Site | Site Score | Dscore | Site volume Å$^3$ | Residue numbers |
|---|---|---|---|---|
| 1 | 1.292 | 1.427 | 304.8 | Chain A & B: 75, 76,77, 78, 128, 129, 130, 13, 132, 134, 165, 166, 167, 168,169, 170, 171 |
| 2 | 1.262 | 1.380 | 266.8 | Chain A: 58,59, 60, 87, 89, 90, 91, 114, 118, 129, 255, 283, 284, 285 Chain B: 93, 95, 97, 110, 111, 112, 134, 136, 137, 138, 160, 162, 164 |
| 3 | 1.109 | 0.983 | 269.9 | Chain B: 55, 56, 59, 88, 90, 111, 113, 115, 117, 133, 135, 137, 163, 175, 177, 181, 192, 194, 196, 212, 216, 218, 231, 232, 233, 244, 262, 266, 267, 268, 269 |
| 4 | 1.098 | 0.962 | 310.4 | Chain A: 39, 40 41, 54, 55, 56, 59, 88, 90, 94, 113, 115, 117, 133, 135, 137, 139, 161, 163, 175, 177, 181, 192, 194, 196, 212, 216, 218, 231, 232, 233, 244, 262, 266, 267, 268, 269, 286, 288 |

**Table S2:** Molecular interaction analysis of small molecules docked at the site-1 StOmpLA dimer

| Ligand name | Docking score | Hydrophobic interactions (Chain) | | | | | Hydrogen bonding (Chain) (atom: ligand atom), Distance Å | | | | | | | | |
|---|---|---|---|---|---|---|---|---|---|---|---|---|---|---|---|
| **Nci97317** | -13.5 | Y76(A, B) | W78(A, B) | P128(A) | F129(B) | P170 (A, B) | | | W78(B) (HE1:O) 1.89 | | | R167(A) (HH11:O HH12:O HH22:O) 1.84, 2.58, 2.66 | S168(A, B) OG:H(A) 1.61 H:O(B) 2.15 H:O(B) 2.59 | | |
| **Alanyl threonine** | -13.2 | Y76(A, B) | W78(A) | | | P170(A) | T75(B)(O:H) 1.99 | N77(B) (H:O) 1.88 | W78(B) (HE1:O) 1.74 | | | R167(A) (HH11:O HH12:O) 2.09, 1.95 | S168(A) OG:H(A) 1.69 | | |
| **Phloretin** | -13.1 | Y76(A, B) | W78(A, B) | P128(A, B) | F129(B) | | | N77(A)(OD1:H) 1.58 | | F129 (B (O:H)2.86 | | | S168(B) OG:H 1.94 | | |
| **VA lactate** | -13.0 | Y76(A, B) | W78(A, B) | | | | | N77(B) (ODE1:H) 2.20 | W78(A) (HE1:O) 1.70 | | | R 167(B, B, A HH12:O, HH12:O) 2.43,1.97, 1.81 | | | |
| **Glycylleucine** | -12.8 | Y76(A, B) | W78(A) | | | P170(B) | | N77(A, A) ODE1:H 2.0,1.97 | W78(A) (HE1:O) 1.90 | | | R167(A, B) (HH11:O(A) HH12:O(B)1.90,2.23 | | | |
| **Aminolevulinic acid** | -12.5 | Y76(A, B) | W78(B) | P128(B) | F129(B) | | T75(B)(O:H) 1.76 | N77(B)(H:O) 2.16 | W78(B) (HE1:O) 1.79 | | | R167(A, B) (HH11:O(A) HH12:O(B) 2.15, 1.74 | | | |
| **Statine** | -12.4 | Y76(A, B) | W78(B) | | | P170(B) | | N77(A, A) ODE1:H 1.79,1.73 | W78(B) (HE1:O) 1.78 | | | R167(A, B) (HH12:O(A) HH11:O(B) 1.84, 1.99 | | | |
| **Nci32977** | -10.3 | Y76(A, B) | W78(A, B) | P128(A, B) | F129(B) | P170(A) | T75(A) (O:H) 1.93; T75(B) (O:H) 1.62 | N77(B) (H:O) 1.76 | | | | R167(B) (HH12:O) 2.01 | S168(B) (O:H) 1.85 | | |
| **Nci36398** | -10.3 | Y76(A, B) | | | | P170 (A, B) | | | | | | | S168(A) (H:O) 1.64, (O:H) 2.34 | D169(B) (O:H) 2.15 | |
| **Nci47582** | -10.3 | Y76(A, B) | W78(A, B) | P128(A) | F129(A) | P170(A) | | N77(B) (O:H) 2.33 | | | | | S168(A) (O:H) 1.87, (O:H) 2.03 | | |
| **Nci14778** | -9.9 | Y76(A, B) | W78(A, B) | P128(A) | | P170 (A, B) | T75(A) (HG1:O) 2.44 | | | | | R167(A) (HH12:O) 2.29; (B) HH11:O) 2.26 | S168(B) (OG:H) 1.85; (H:O) 2.26 | | T171(B) (OG1:H) 1.93 |
| **Nci19775** | -9.9 | Y76(A, B) | W78(A, B) | P128(A) | F129(A) | P170 (A, B) | T75(B) (O:H) 1.98 | N77(A) (H:O) 2.65; (B) (OD1:H) 1.87, (OD1:H) 1.84 | | | | | S168(A) (OG:H) 2.43 | | |
| **Nci2819** | -9.8 | Y76(A, B) | W78(A) | | | P170 (A, B) | T75(B) (HG1:O) 2.57, (O:H) 1.90 | | | | | | S168(A) (H:O) 2.46, (O:H) 1.97; (B) (H:O) 2.07 | D169(B) (O:H) 2.30 | |
| **Sulphomeoxayzole** | -6.8 | Y76(A, B) | W78(A, B) | P128(A, B) | F129(A, B) | P170(B) | | | | F129 (A) (O:H)2.09, (B) (O:H) 2.11 | G166(B) (O:H) 2.73 | R167(A) (HH22: N) 2.68, (B) (HH22: N) 2.30 | S168(A) (H:O) 2.03, (B) (H:O) 1.84 | | |

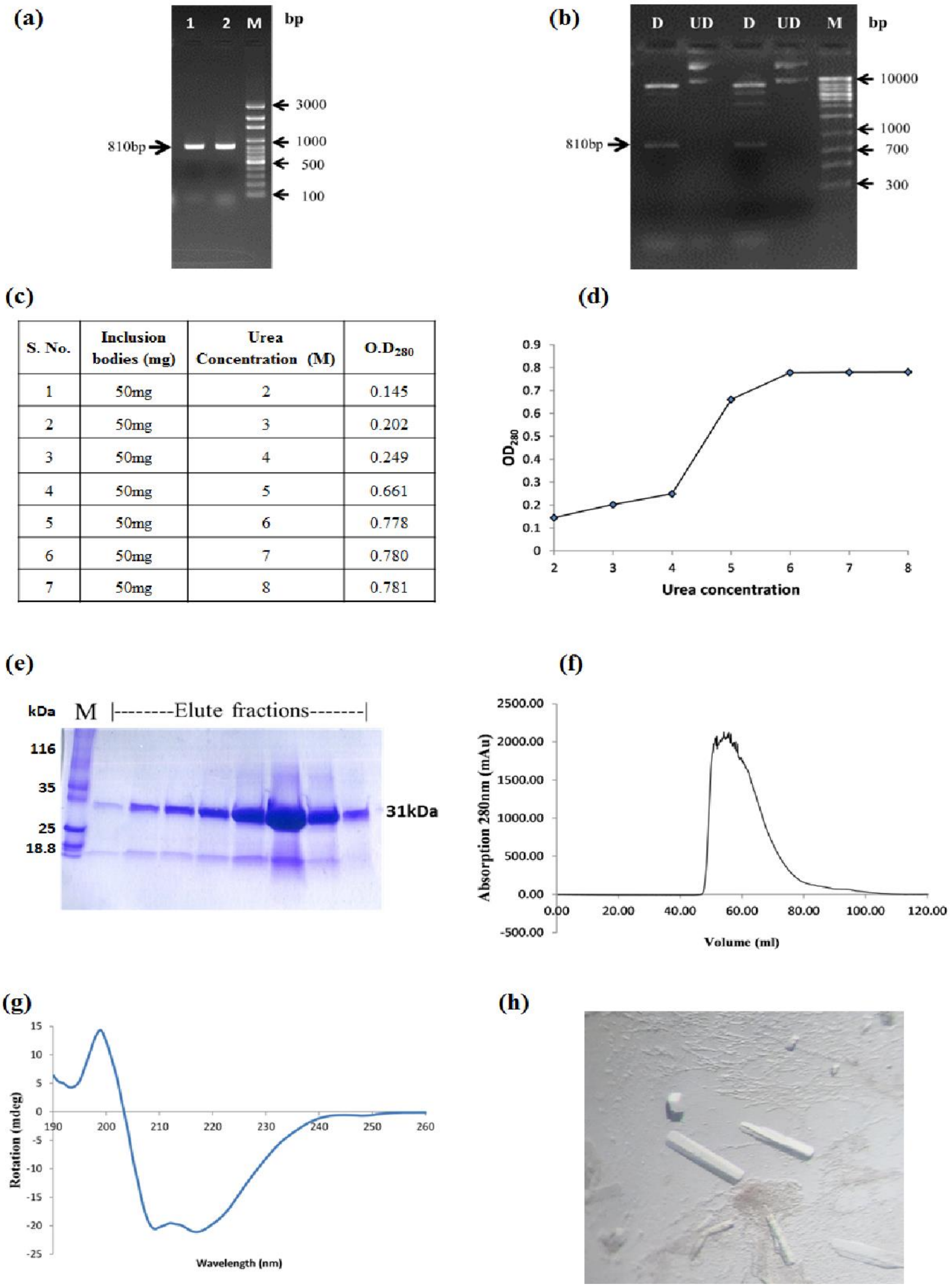


| S. No. | Inclusion bodies (mg) | Urea Concentration (M) | O.D$_{280}$ |
|---|---|---|---|
| 1 | 50mg | 2 | 0.145 |
| 2 | 50mg | 3 | 0.202 |
| 3 | 50mg | 4 | 0.249 |
| 4 | 50mg | 5 | 0.661 |
| 5 | 50mg | 6 | 0.778 |
| 6 | 50mg | 7 | 0.780 |
| 7 | 50mg | 8 | 0.781 |

**Figure S1.** Cloning, refolding, and crystallization of StOmpLA. **(a)** PCR amplification *of S. typhi*, *pldA* gene. Lane 1, 2 show PCR products and lane 3 is shows 1 kb ladder. **(b)** Double digestion of *pldA*-pET30b (810bp insert) construct with *Nde*I and *Bam*HI; D: Digested plasmid DNA, UD: Undigested plasmid DNA, M: 100 bp DNA ladder. **(c)** and **(d)** summarize optimisation conditions for refolding of OmpLA from inclusion bodies using UV spectrophotometer. (**e**) Denaturing and reducing SDS-PAGE gel showing eluted fractions from Superdex S200 column, M: Marker, and (**f**) corresponding chromatogram. (**g**) CD spectrum for purified *S. typhi* OmpLA in $C_{12}E_9$. (**h**) OmpLA crystals obtained in 0.1 M sodium iodide, 0.1 M sodium phosphate (pH 7.0), and 33% v/v polyethylene glycol 300.

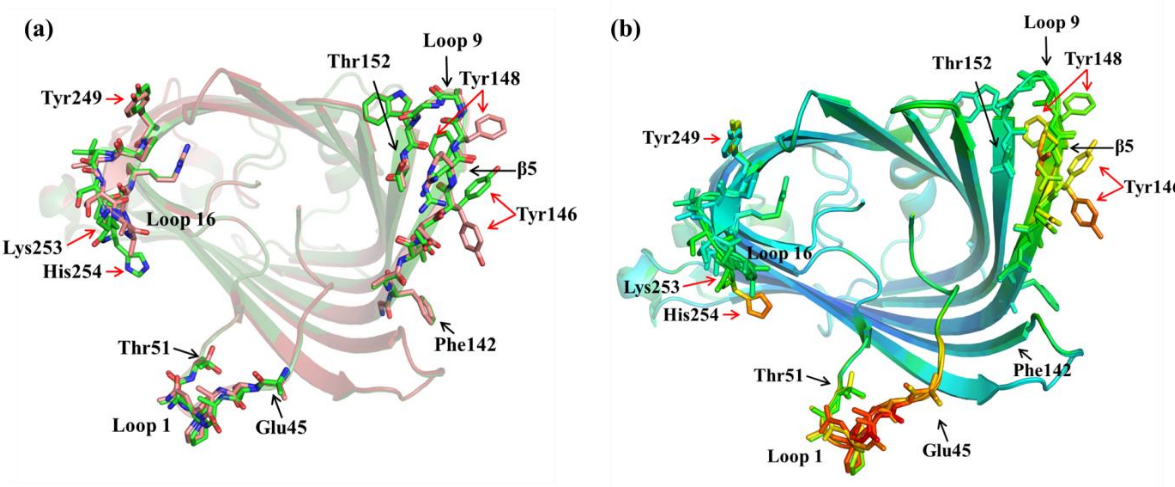


**Figure S2.** Structural superposition of StOmpLA monomeric subunits of the functional dimer. (**a**) Chain A (pink) and chain B (green) of StOmpLA with structural differences marked with red arrows, and **(b)** B-factor differences in the loops and β-strands in both chains, marked by red arrows.

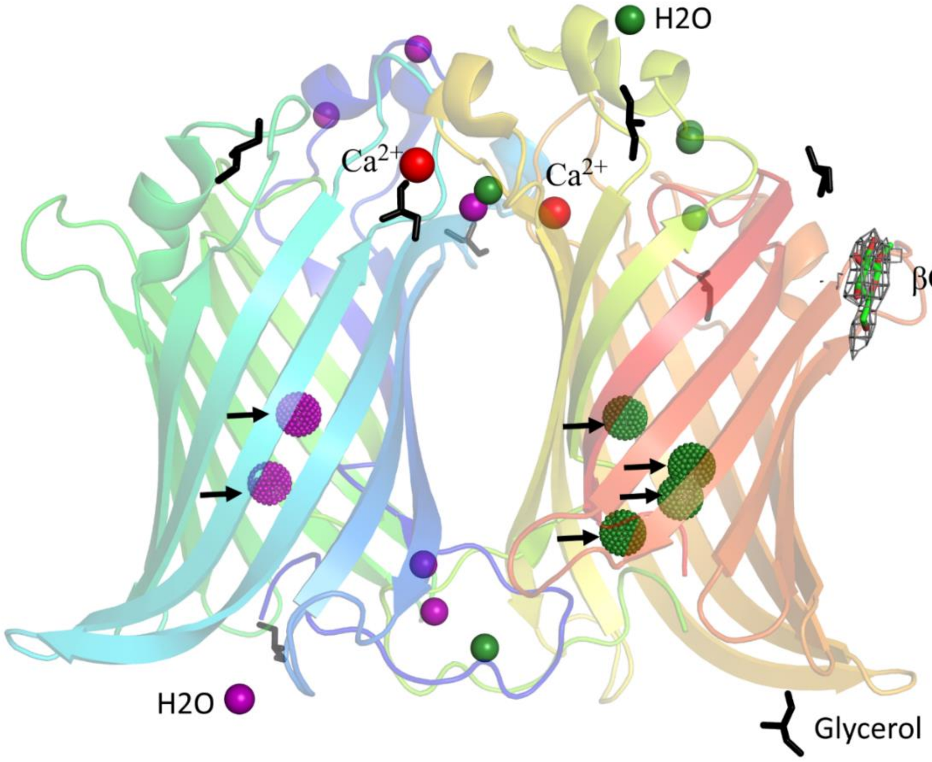


**Figure S3**. OmpLA dimer showing water molecules as spheres. Waters coordinated by chain A and B are color coded in purple and forest green, respectively. Water molecules lining the channel like interior of the barrel are represented as dots (marked by arrows), two calcium atoms in red, and glycerol molecules as black sticks. β-OG bound to chain B is shown with 2Fo-Fc map contoured at 1 sigma.

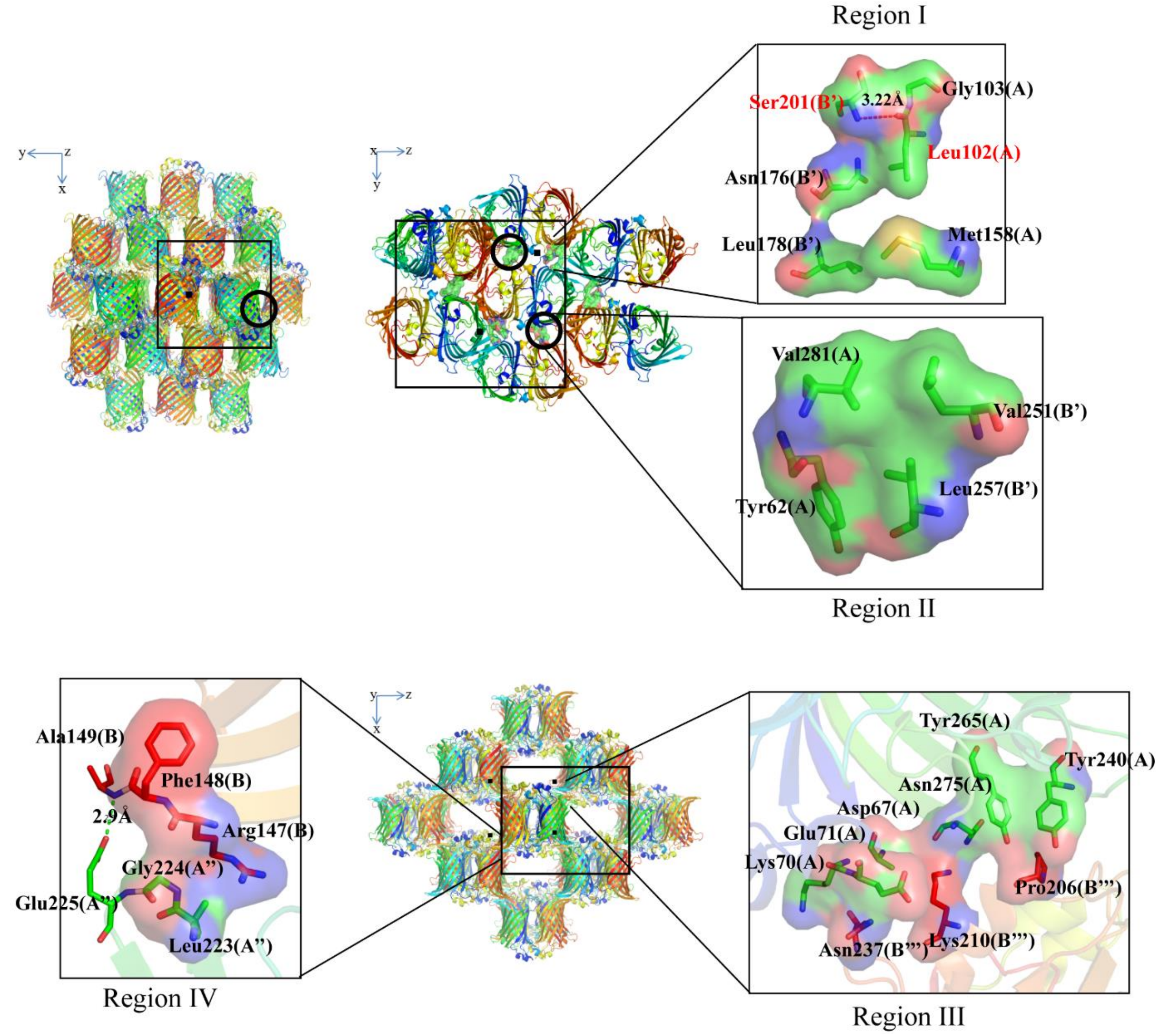


**Figure S4.** Crystal packing in StOmpLA**.** Type II packing showing various crystal contacts in the unit cell of StOmpLA. YZ plane shows crystal contacts formed by two hydrophobic patches (region I and II) on both the highly convex sides of the protein. XZ plane shows contacts formed through region IV hydrophobic patch as well as region III.

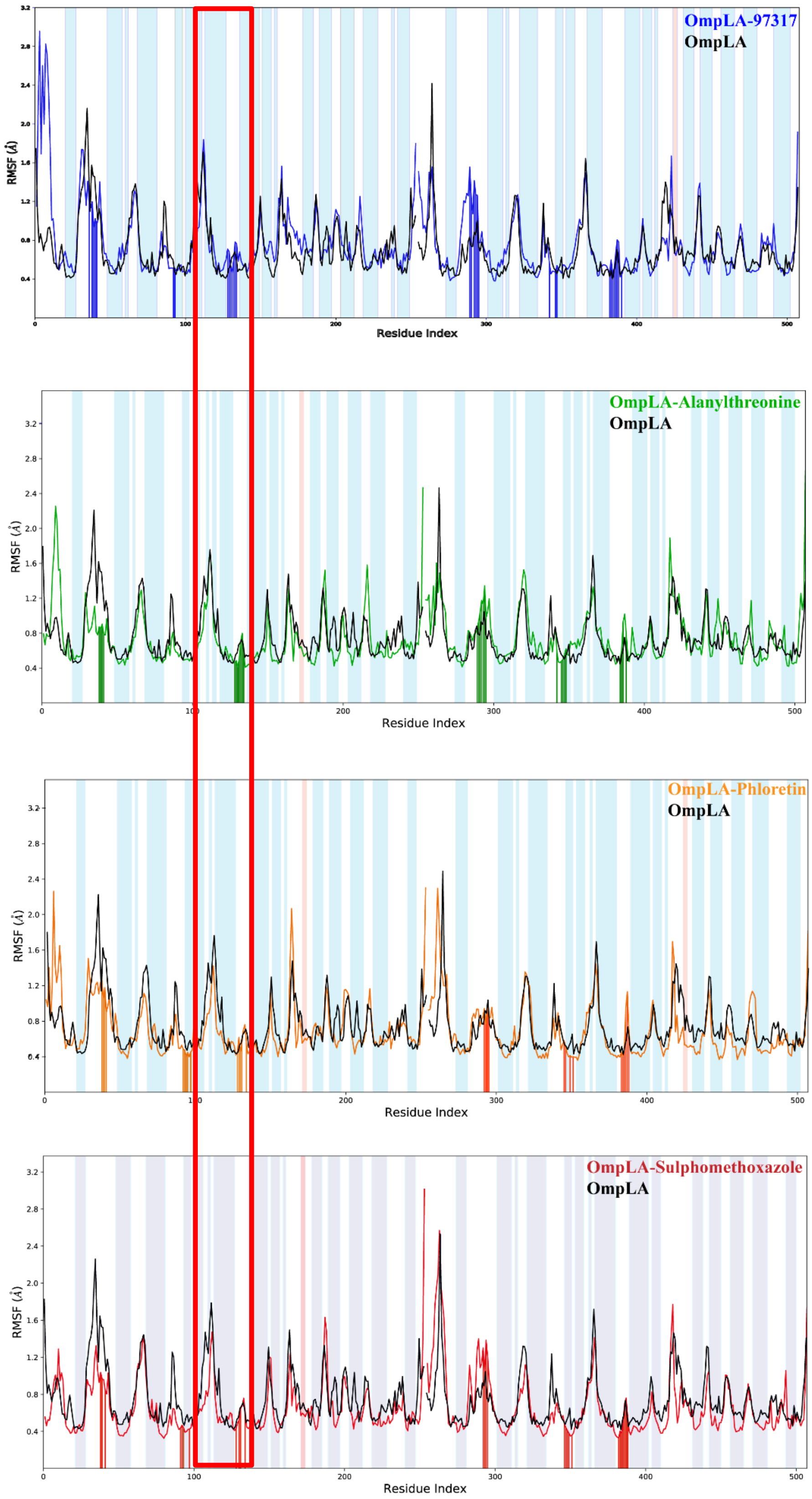


**Figure S5**. RMSF plot for three top hits and sulfamethoxazole throughout 100ns simulation and the red box demarcates the extracellular part of barrel covering region between loops L3 and L5.

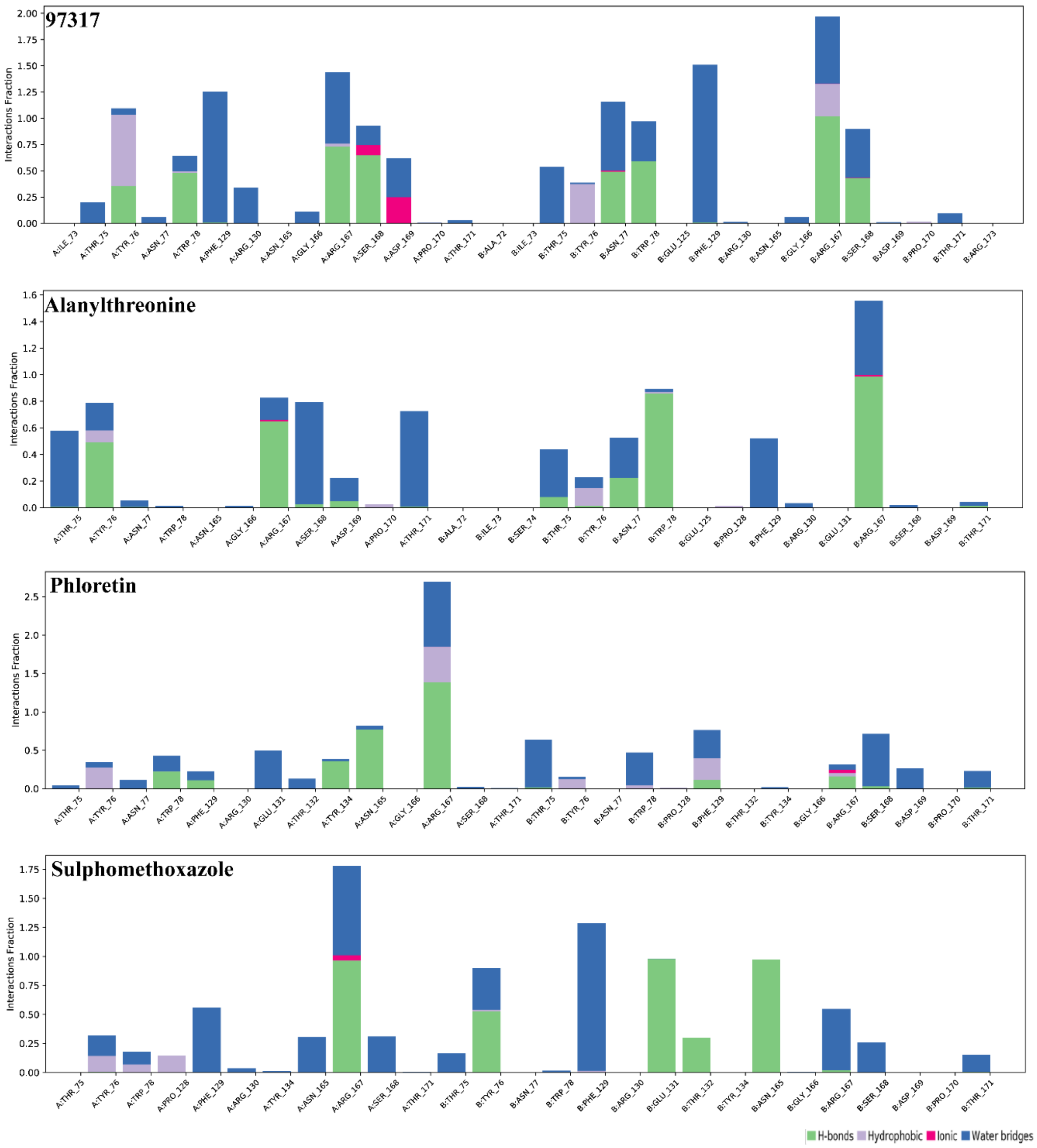


**Figure S6**. Molecular interaction analysis of OMPLA with top ranked hits and sulfamethoxazole throughout 100ns simulation.

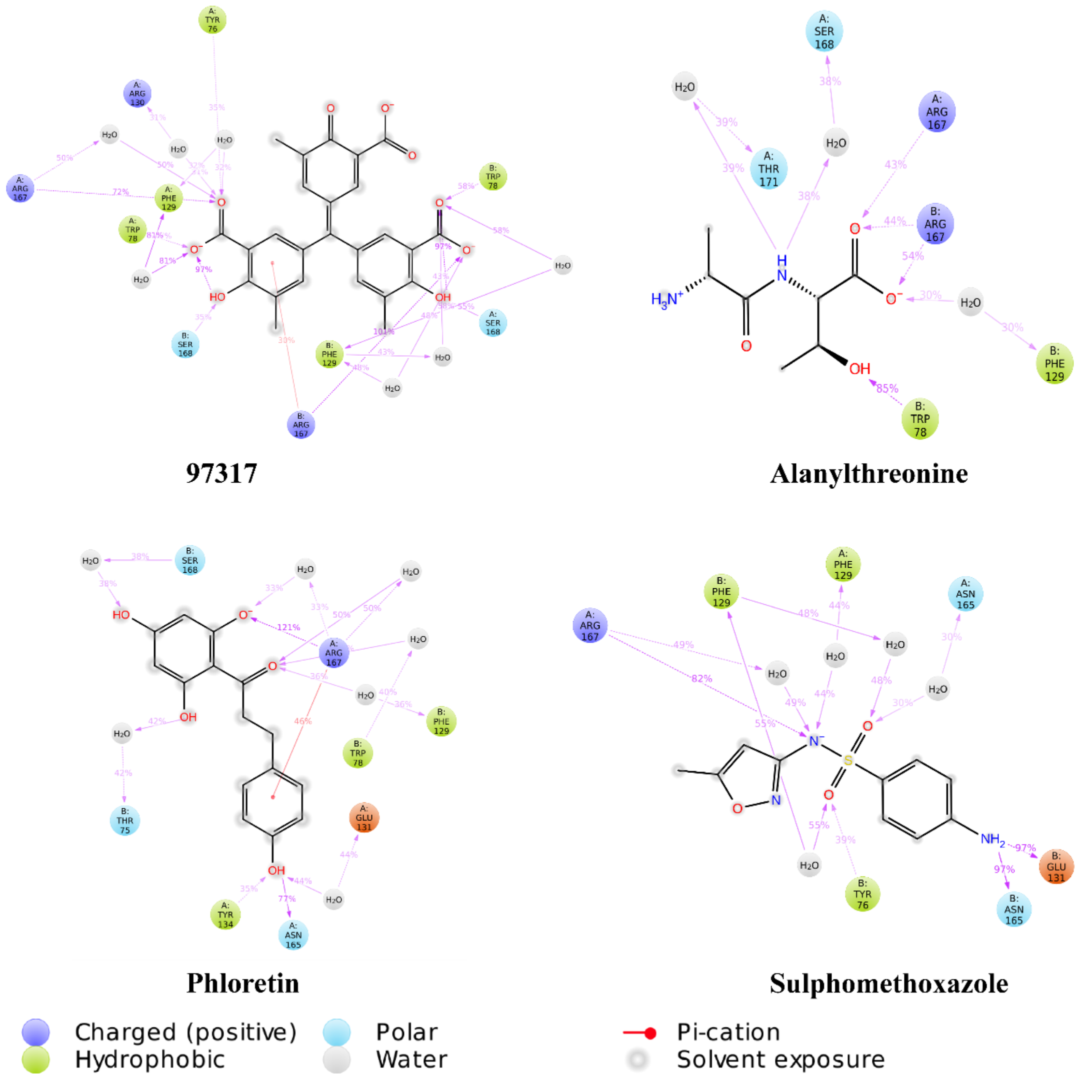


**Figure S7**. Molecular interaction analysis of OmpLA with top ranked hits and sulfamethoxazole showing 2D view of interactions with 30%-time occupancy throughout 100 ns simulation.

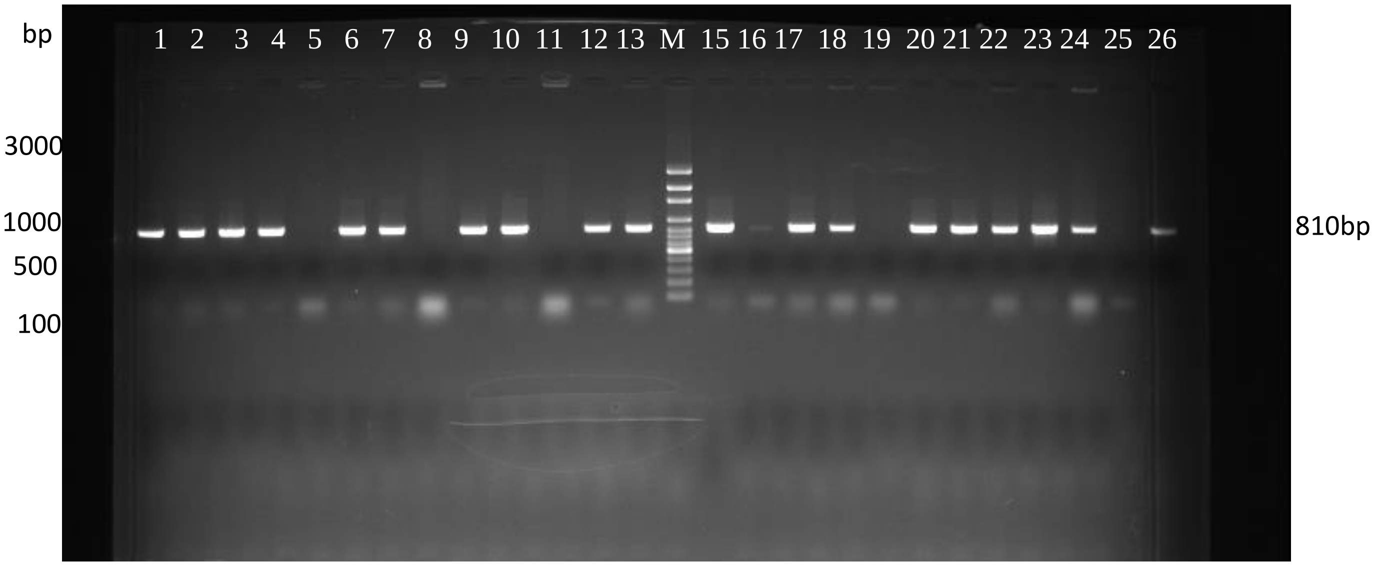


Uncropped agarose gel used in Fig. S1a. Gradient PCR of *S. typhi*, *pldA* gene. Lane 1- 4, 6, 7, 9, 10, 12, 13, 15-18, 20-24, 26 were loaded with PCR products and lane "M" loaded with 100 bp ladder. The expected size of the PCR product is 810 bp.

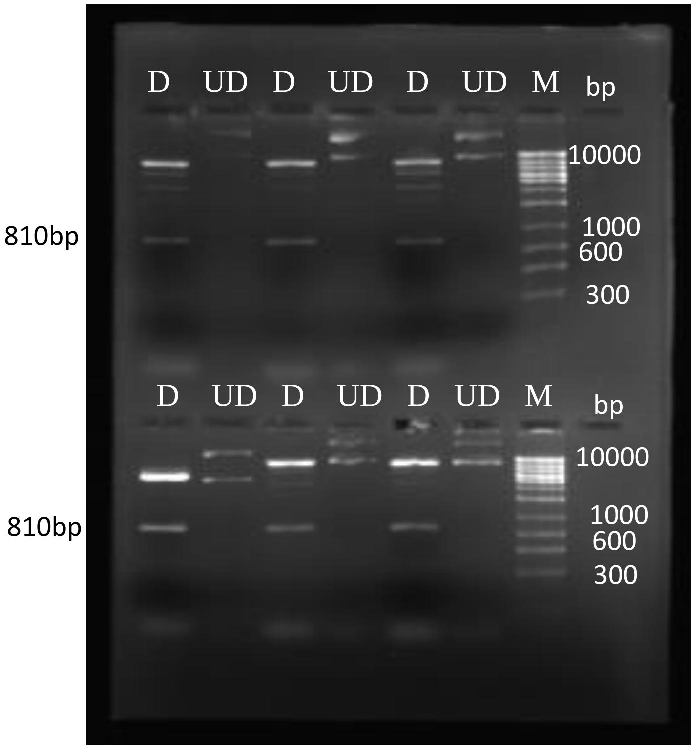


Uncropped agarose gel used in Fig. S1b. Double digestion of cloned into *pldA*-pET30b (810bp insert) with *Nde*I and *Bam*HI. D: Double digested plasmid DNA, UD: Undigested plasmid DNA, M: 1kb DNA ladder.

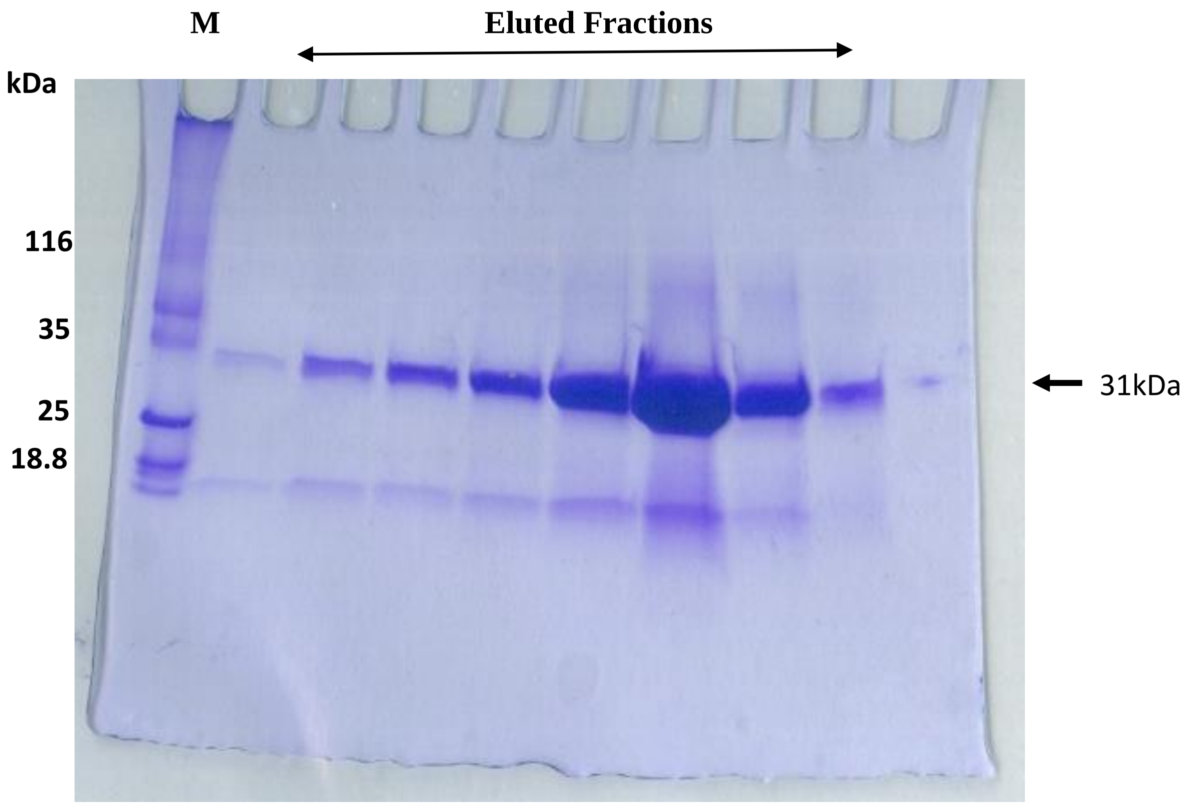


Uncropped SDS-PAGE gel used in Fig. S1e. Denaturing and reducing 4-20% gradient SDS-PAGE gel shows the eluted fractions from Superdex S200 column, M: Standard molecular weight markers.
